# Altered microRNA Expression Correlates with Reduced TLR2/4-Dependent Periodontal Inflammation and Bone Resorption Induced by Polymicrobial Infection

**DOI:** 10.1101/2025.01.10.632435

**Authors:** Syam Jeepipalli, Parvathi Gurusamy, Ana Rafaela Luz Martins, Eduardo Colella, Sandhya R Nadakuditi, Tushar Desaraju, Ashitha Yada, Jennifer Onime, John William, Indraneel Bhattacharyya, Edward K. L. Chan, L. Kesavalu

## Abstract

Periodontitis (PD) is a polymicrobial dysbiotic immuno-inflammatory disease. Toll-like receptors (TLRs) are present on gingival epithelial cells and recognize pathogen-associated molecular patterns (PAMPs) on pathogenic bacteria, induce the secretion of proinflammatory cytokines, and initiate innate and adaptive antigen-specific immune responses to eradicate the invading microbes. Since PD is a chronic inflammatory disease, TLR2/TLR4 plays a vital role in disease pathogenesis and maintaining the periodontium during health. Many factors modulate the TLR-mediated signaling pathway, including specific miRNAs. The present study was designed to characterize the function of TLR2/4 signaling to the miRNA profile after polybacterial infection with *Streptococcus gordonii, Fusobacterium nucleatum, Porphyromonas gingivalis, Treponema denticola,* and *Tannerella forsythia* in C57BL6/J wild-type, TLR2^−/−^, and TLR4^−/−^ mice (n=16/group) using RT-qPCR. The selection of 15 dominant miRNAs for RT-qPCR analysis was based on prior NanoString global miRNA expression profiling in response to polymicrobial and monobacterial infection. Polybacterial infections established gingival colonization in wild-type, TLR2^−/−^ and TLR4^−/−^ mice with induction of bacterial-specific IgG. A significant reduction in alveolar bone resorption (ABR) and gingival inflammation was observed in the mandibles of TLR2/4^−/−^ mice compared to C57BL6/J wild-type mice (*p*<0.0001). Periodontal bacteria disseminated from gingival tissue to the multiple organs in wild-type and TLR2^−/−^ mice (heart, lungs, brain, kidney) and limited to heart (*F. nucleatum*), lungs (*P. gingivalis*), kidney (*T. forsythia*) in TLR4^−/−^ mice. The diagnostic potential of miRNAs was assessed by receiver operating characteristic (ROC) curves. Among 15 miRNAs, three were upregulated in C57BL6/J wild-type mice, two in TLR2^−/−^, and seven in TLR4^−/−^ mice. Notably, the anti-inflammatory miR-146a-5p was consistently upregulated in all the mice. Additionally, miR-15a-5p was upregulated in wild-type and TLR2^−/−^ mice. let-7c-5p was upregulated in TLR4^−/−^ mice and downregulated in the wild-type mice. Multi-species oral bacterial infection alters the TLR2/4 signaling pathways by modulating the expression of several potential biomarker miRNAs in periodontium.

**IMPORTANCE:** Periodontitis is the most prevalent chronic immuno-infectious multispecies dysbiotic disease of the oral cavity. The Toll-like receptors (TLR) provide the first line of defense, one of the best-characterized pathogens-detection systems and play a vital role in recognizing multiple microbial products. Multispecies infection with periodontal bacteria *S. gordonii, F. nucleatum, P. gingivalis, T. denticola,* and *T. forsythia* induced gingival inflammation, alveolar bone resorption (ABR) and miRNA expression in the C57BL6/J wild-type mice and whereas infection did not increase significant ABR in the TLR2/4 deficient mice. Among the 15 miRNAs investigated, miR-146a**-**5p, miR-15a-5p were upregulated in wild-type and TLR2^−/−^ mice and miR-146a-5p, miR-30c-5p, let-7c-5p were upregulated in the TLR4^−/−^ mice compared to sham-infected controls. Notably, inflammatory miRNA miR-146a-5p was upregulated uniquely among the three different infection groups. The upregulated miRNAs (miR-146a, miR-15-a-5p, let-7c-5p) and downregulated miRNAs could be markers for TLRs-mediated induction of periodontitis.

## INTRODUCTION

Periodontitis (PD) is an immuno-inflammatory polymicrobial dysbiotic disease characterized by complex subgingival plaques and leads to the inflammatory destruction of the supporting tissues, including gingival tissue, periodontal ligament, and alveolar bone. Several studies report that *P. gingivalis*, *T. forsythia*, *T. denticola,* and *F. nucleatum* (co-aggregating bacteria) are more frequently identified, synergistically interact with each other, and in higher numbers in adult periodontitis as compared to healthy individuals, and are also positively correlated with pocket depth and bleeding on probing, measures of periodontal tissue destruction [1].

MicroRNAs (miRNAs) are small noncoding RNA molecules that play critical role in regulating gene expression by directly binding to their 3′ untranslated regions [2,3]. Certain miRNAs are implicated in the pathogenesis of PD [4–7]. The periodontal miRNAs expressed in the polybacterial infection (miR-375, miR-690, miR-148a, mmu-let-7a-5p) are not identical to those in monobacterial infection such as *P. gingivalis* (miR-133a, miR-22)*, T. denticola* (miR-133a, miR-378, miR-34b-5p), *T. forsythia* (miR-1902, miR-720)*, F. nucleatum* (miR-361, miR-323-3p)*, S. gordonii* (miR-135a, miR-720) [8–13].

Clinical studies have shown that mRNA expression of TLR2 and TLR4 is significantly elevated in gingivitis [14] and in tissues affected by severe periodontitis [15,16]. These results showed that TLR2 and 4 played a major role in pathogenesis. It is known, for example, lipoprotein from *P. gingivalis* [17] and other bacteria are known to activate TLR2 [18] while *P. gingivalis* LPS activates TLR4 [19–21]. Also, our report focusing on atherosclerosis in this mouse model shows that polybacterial infections have established gingival colonization and induction of a pathogen-specific IgG immune response in TLR2^−/−^ and TLR4^−/−^ mice after several weeks of chronic gingival infection [22]. TLR2/4 deficiency dampened ABR and intrabony defects, indicating TLR2/4 central role in polymicrobial infection-induced periodontitis [22]. Thus, it is interesting to define the relative contribution of TLR2/4 and the function of their associated dominant miRNAs to the pathogenesis in our polymicrobial PD model. TLR2 is detected in gingival pocket epithelia, gingival fibroblasts, neutrophils, cementum, periodontal ligament fibroblasts (PLFs), osteoclasts, and tissue dendritic cells whereas TLR4 is predominantly detected in the gingival epithelium, fibroblasts, cementum, PLFs, dendritic cells, osteoblasts and endothelium [23,24]. TLRs provide the first line of defense and one of the best-characterized pathogens-detection systems. They play a vital role in recognizing multiple microbial products, including LPS, Lipoproteins, Peptidoglycan, Lipoteichoic acid, and fimbriae. Activation of TLRs triggers the release of proinflammatory mediators such as TNFα, IL-1β, and IL-8 through the induction of transcription factors NF*k*B (nuclear factor kappa-light-chain-enhancer of activated B cells); this signaling pathway is essential for initiating immune response and driving gingival inflammation and resulting adaptive immune responses. TLR2 recognizes peptidoglycans, lipoproteins, and lipoarabinomannans from Gram-positive bacteria, and TLR4 is activated by the Gram-negative bacterial component LPS. In addition, several miRNAs can bind to TLRs and initiate adaptive immune responses by inducing immune and inflammatory gene expression. TLRs (2,4) are expressed both on immune cells (dendritic cells, macrophages, B cells, natural killer cells) and in periodontal tissues [25], interact with pathogen-extracellular stimuli (MAMPs) [26], and play a central role in gene expressions, inflammation, and different stages of the periodontitis [15,27].

Periodontal bacteria surface antigens in gingivitis and periodontitis interact with TLRs and play an essential role in developing adaptive immunity [28,29]. Reciprocal extracellular miRNA vesicle communication between different cell populations facilitates the host response [30]. In our previously published study, mice deficient in TLR2 and 4 receptors affected both periodontitis and atherosclerosis [22,31] to polymicrobial infections. Identifying the miRNA markers at the early stages of PD is essential to initiate timely interventions that prevent disease complications and preserve oral health. Diagnostic miRNA discovery from the preclinical studies could help clinicians identify the disease severity in human periodontitis, e.g., miR-146a-5p [32,33]. The high expression of miR-146a was observed in many inflammatory diseases, such as osteoarthritis and rheumatoid arthritis [34].

The preclinical *in vivo* studies involving five different bacteria (*S. gordonii, F. nucleatum, P. gingivalis, T. denticola, T. forsythia*) have revealed specific differentially expressed (DE) miRNAs in each infection that was analyzed using NanoString nCounter technology [8–11,13]. These miRNAs exhibit a unique range of expression patterns during polymicrobial infections [8] as well as five distinct monoinfection (*S. gordonii, P. gingivalis, F. nucleatum, T. denticola, T. forsythia*) [9,11,13] with highest upregulated and downregulated miRNAs with high fold chain, 15 miRNAs were selected for investigation of its presence in C57BL6/J wild-type (hear after termed as wild-type), TLR2^−/−^, and TLR4^−/−^ gene knockout mice (Table 2) mandibles to polymicrobial infection. The 15 selected miRNAs are mmu-let-7c-5p, miR-15a-5p, miR-22-5p, miR-30c-5p, miR-34b-5p, miR-133a-3p, miR-146a-5p, miR-323-3p, miR-339-5p, miR-375-3p, miR-361-5p, miR-423-5p, miR-720, miR-155-5p, and miR-132-3p. Reverse-transcription quantitative polymerase chain reaction (RT-qPCR) is an effective method for studying gene expression and measuring the expression level of target gene expressions. In the present study, we aimed to investigate the critical function of 15 preselected dominant periodontal miRNAs using RT-qPCR in polymicrobial infected (mimicking the human oral microbiota ecological colonization) wild-type, TLR2^−/−^ and TLR4^−/−^ male and female mice mandibles.

## MATERIALS AND METHODS

### Experimental mice and housing conditions

This experiment used wild-type, TLR2^−/−^, and TLR4^−/−^ mice purchased from Jackson Laboratory (Bar Harbor, ME, USA). Upon arrival, the mice were allowed a week of acclimation before initiating the infection and sham infection. At the time of infection, we used nine-week-old mice for this study. Throughout the study, mice were housed in a controlled environment with 12 hours of dark/light cycles and maintained at a consistent temperature. Mice had access to standard chow and water ad libitum. All animal procedures were conducted in accordance with ethical guidelines approved by the University of Florida Institutional Animal Care and Use Committee under protocol number 202200000223. Mice were randomly assigned to the infection and sham infection group for each genotype (wild-type, TLR2^−/−^ and TLR4^−/−^).

### Grouping the mice

Mice were divided into polybacterial infection groups and sham infection groups. Each group consists of sixteen mice (8 male and eight female). The sample size was determined based on previous studies from Aravindraja et al. [8,9,11,13]. Mice in group I (wild-type mice), III (TLR2^−/−^ mice), and V (TLR4^−/−^ mice) underwent sequential polymicrobial infection cycles (ICs). These infection cycles followed a specific order: *S. gordonii* (*Sg*, 2 ICs)*, F. nucleatum* (*Fn*, 2 ICs), and later with *P. gingivalis* (*Pg*), *T. denticola* (*Td*), and *T. forsythia* (*Tf*) (5 ICs). Each infection cycle consisted of four days of intraoral bacterial infection in a week, using 2.5×10^8^ cells of each bacteria species suspended in reduced transport fluid (RTF)+6% CMC [8], while sham-infected Group II (wild-type), Group IV (TLR2^−/−^), and Group VI (TLR4^−/−^) mice were mock-infected with RTF+4% CMC only (Figure 1A).

**FIGURE 1.**
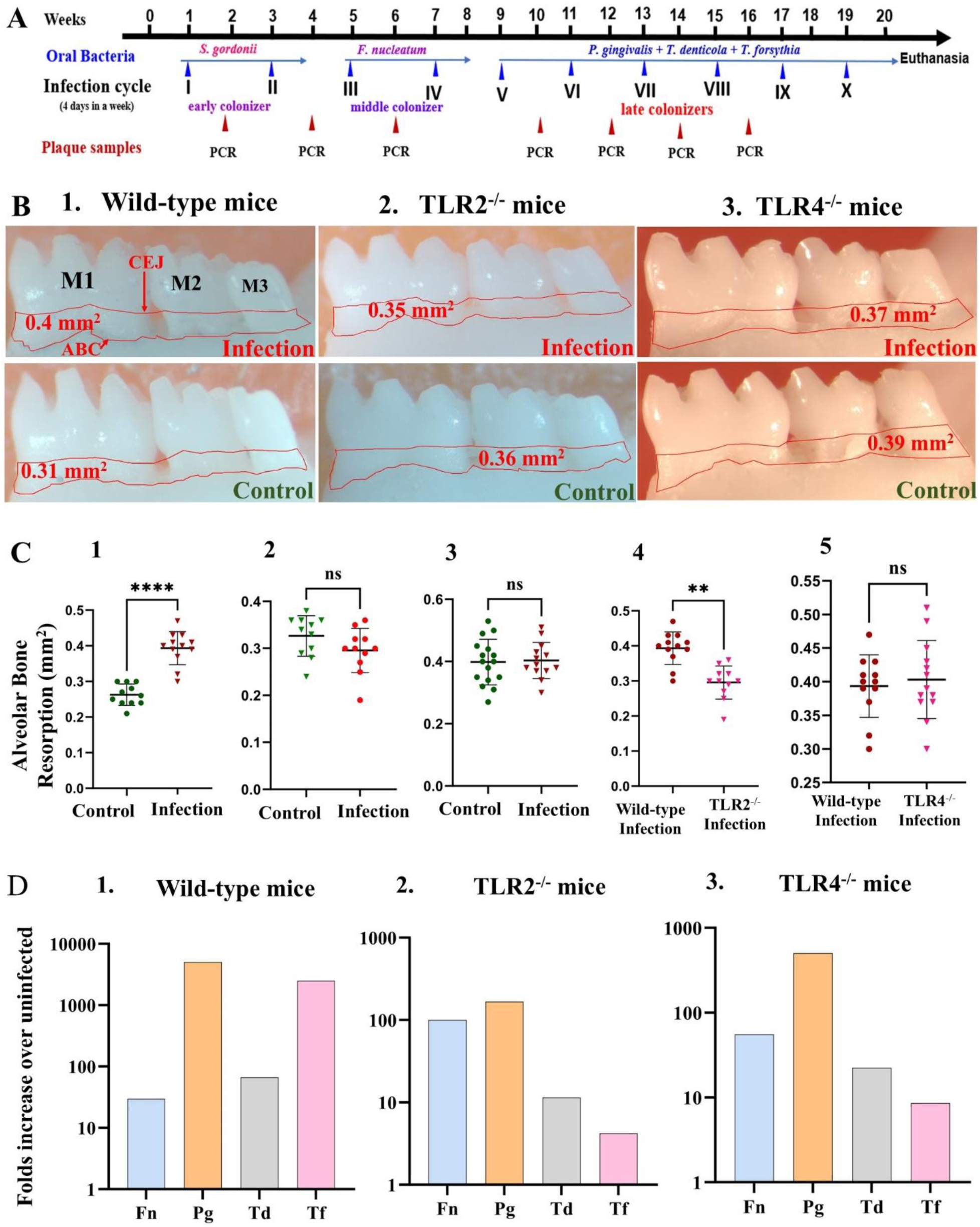
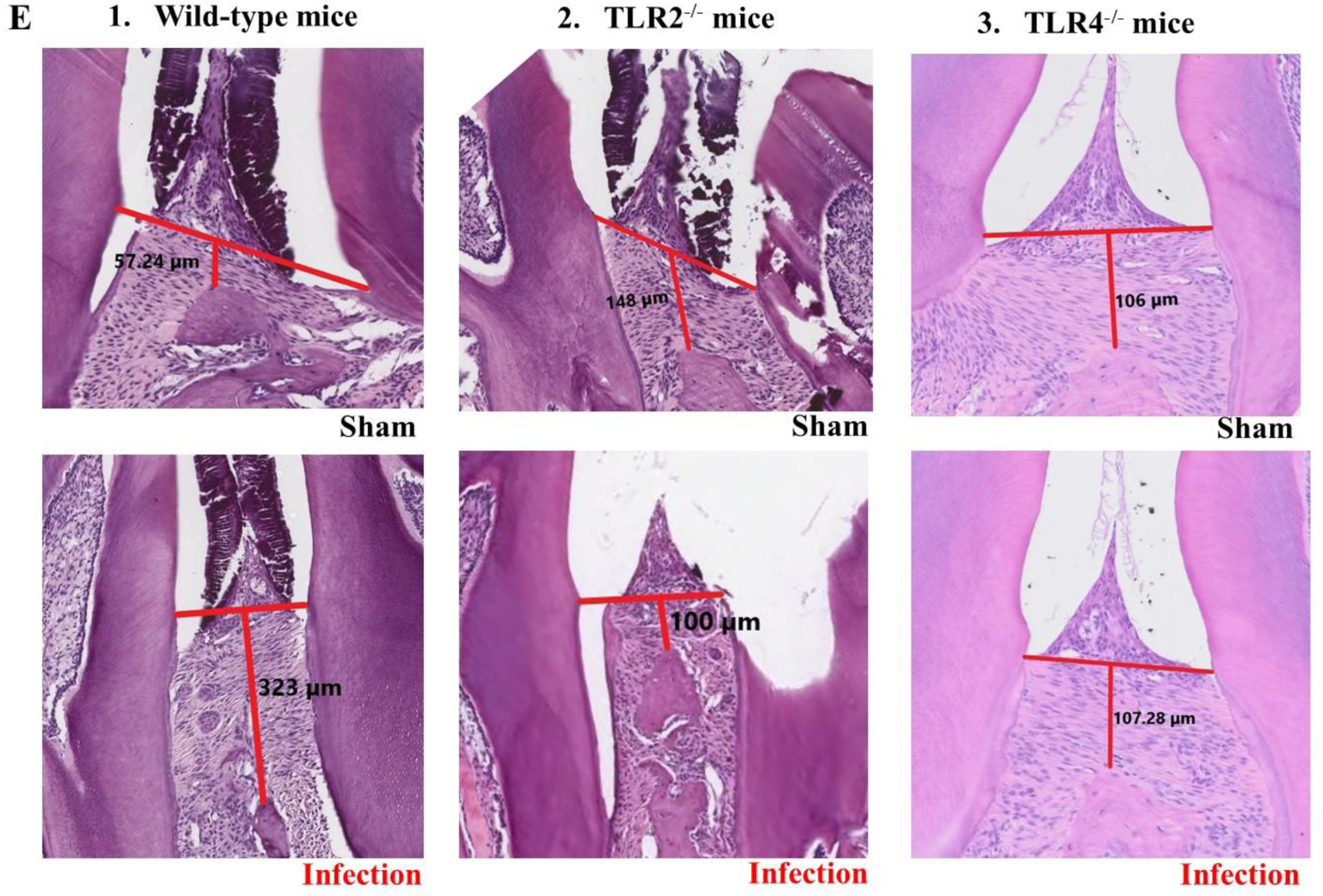
Intraoral Ecological Time-sequential Polymicrobial Periodontal Infection (ETSPPI) results in alveolar bone resorption. (**A**) Schematic diagram of the experimental design depicting the ETSPPI infection (4 days per week on every alternate week), plaque sampling for PCR, and euthanasia. (**B**) Representative images showing horizontal ABR (mandible lingual view) of the polybacterial infected and sham-infected mice with the area of ABR outlined from the alveolar bone crest (ABC) to the cementoenamel junction (CEJ). (**C**) Morphometric analysis of the mandible and maxillary ABR in mice. A significant increase in ABR was observed in the mandible lingual (p < 0.0001) and maxilla buccal (p < 0.01) in the wild-type mice. Polymicrobial infection did not show ABR in the mandible lingual, maxilla palatal, and maxilla buccal sides in the TLR2^−/−^ mice and TLR4^−/−^ mice relative to their uninfected control mice. **(D)** Bacteria-specific IgG antibody analysis. Ordinary two-way ANOVA). Data points and error bars are mean± SEM (*n* = 16). (**E)** Representative H&E-stained histological mandible tissue sections from polymicrobial infected and sham-infected wild-type (1E left), TLR2^−/−^ (1E middle), and TLR4^−/−^ mice (1E right) (n=5) showing epithelial apical migration and infiltration of inflammatory cells. Brackets indicate locations of gingival junctional epithelial (JE) migration. Arrowheads indicate the cementoenamel junction, where the gingival junctional epithelium naturally ends. Bar =100 µm. Images are shown at 40X magnification.

### Bacterial culture and oral administration

In this study, the following bacterial strains were used: *S. gordonii* DL1, *F. nucleatum* ATCC 49256, *P. gingivalis* ATCC 53977, *T. denticola* ATCC 35404, and *T. forsythia* ATCC 43037. Standard protocol for bacterial growth culture, harvesting, and cell counting procedures were followed as described by Aravindraja *et al*. [8,22]. Before bacterial infection, mice were allowed to consume Kanamycin water (500 mg/2L) for 3 days and rinsed the oral cavity with 0.12% chlorohexidine gluconate to further reduce bacterial growth [8].

### Molecular detection of bacteria in the gingival swabs and systemic organs

Gingival swabs from each mouse were collected after 4^th^ day of an infection cycle using a sterile cotton swab, then suspended Tris-EDTA (TE) buffer and subjected to colony polymerase chain reaction (PCR). Testing was performed in Bio-Rad thermal cycler (Bio-Rad, Hercules, CA, USA) with the test reaction components such as master mix (NEB, Ipswich, MA, USA), 16s rRNA specific forward primer (FP), reverse primer (RP), and the DNA source (oral swabs). Details of each bacterial primer are shown in Table 1. DNA extracted from pure bacterial culture was considered a positive control, and sterile PCR-grade water was a negative control. PCR products were run in 1% agarose gel electrophoresis and visualized using UVP GelStudio Touch Imaging System (Analytik Jena US LLC, CA, USA) [8,22]. Bacterial genomic DNA (gDNA) from the organs was harvested following the standard protocol described in the Qiagen Dneasy Blood and Tissue kit (Qiagen, Germantown, MD, USA). The PCR test was performed as described above.

**TABLE 1.**
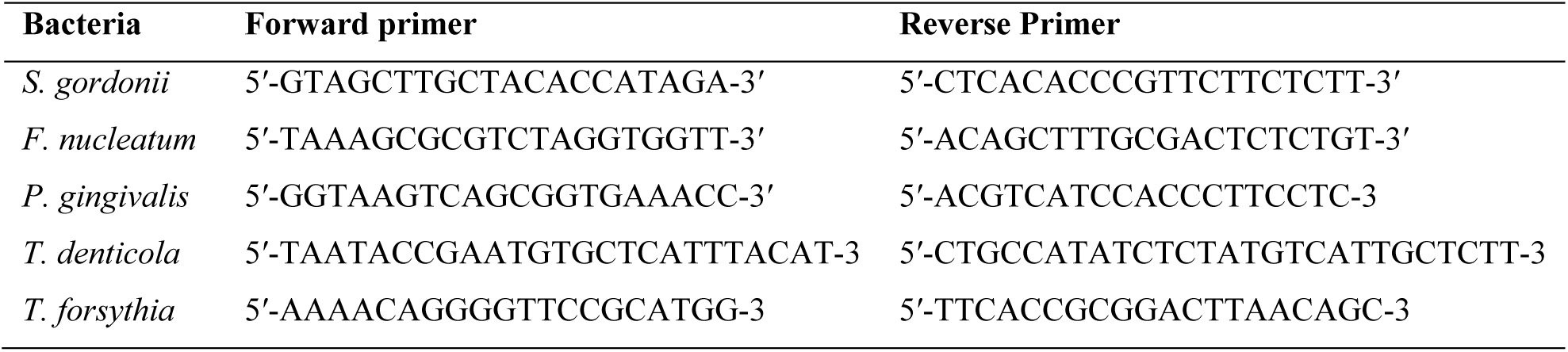
List of 16s rRNA-specific primer sequences for periodontal bacteria.

### Euthanasia and sample collection

Mice were euthanized a week after the 10^th^ infection cycle following the carbon dioxide (CO_2_) inhalation procedure. Blood was drawn through cardiac puncture procedure, and the serum was isolated and stored at –20°C to measure immunoglobulin-G (IgG) concentration against each bacterium. Additionally, the distal organs (heart, liver, kidney, spleen, lungs, and brain) were harvested and stored at –80°C for further examination. The left maxilla and mandibles were excised and preserved in RNA*l*ater at –80°C for the total RNA extraction. The right maxilla and mandibles were excised and processed for alveolar bone resorption (ABR) morphometry analysis.

### Serum antibody analysis

Serum isolated from the wild-type mice, TLR2^−/−^ mice, and TLR4^−/−^ mice was used to analyze the levels of periodontal bacteria (*Sg, Fn, Pg, Td,* and *Tf*) surface antigen-specific IgG was determined separately by enzyme-linked immunosorbent assay (ELISA) as described Aravindraja et al. [8,35]. Briefly, the formaldehyde-killed bacteria such as *F. nucleatum*, *P. gingivalis*, *T. denticola*, and *T. forsythia* were used for coating antigens to the ELISA plate wells, and diluted serum (100µL) of each mouse was added to each well in triplicates. Goat anti-mouse-IgG alkaline phosphatase (Sigma Aldrich, St. Louis, MO, USA) was added to each reaction well, incubated, and color developed with *p*-Nitrophenyl phosphate (Sigma Aldrich, St. Louis, MO, USA). Color development was stopped by adding 3M NaOH, and the yellow color intensity was measured at OD_405 nm_ using an Epoch microplate spectrophotometer and analyzed in Gen5 software (BioTek, Winooski, VT, USA). The infected mice serum antibody was quantified using a gravimetric standard curve (Sigma Aldrich).

### Histology of gingival tissue

The right mandibles from the polymicrobial-infected and sham-infected (wild-type, TLR2^−/−^, and TLR4^−/−^) group’s mice (n=3) were decalcified in Immunocal (Decal Chemical, Tallman, NY) for 28 days at 4°C. The decalcified mandibles were embedded in paraffin blocks and sectioned (4 μm) along the mesiodistal plane. Sections were stained with hematoxylin and eosin (H&E) for histological analysis, and slides were scanned with a ScanScope CS system (Aperio, Vista, CA). The digitally scanned slides were viewed at ×200 magnification objective lens, ScanScope™ XT (Aperio Technologies, Inc., Vista, CA, USA), and analyzed using the ImageScope program [36,37]. Evidence of inflammation, such as apical junctional epithelial (JE) migration, elongation of rete ridges, resorption of alveolar bone, epithelial hyperplasia, and epithelial edema was determined [38,39].

### Measurement of alveolar bone resorption (ABR)

The impact of polymicrobial infection on alveolar bone was assessed following the histomorphometry procedure [8,22]. After euthanasia, the excised mandibles and maxilla were placed in a beaker and autoclaved to remove the soft tissue. The defleshed bone specimen was cleaned in 3% H_2_O_2_ solution and air-dried. Two-dimensional imaging of the bone was captured under a 10×stereo dissecting microscope (SteReo Discovery V8, Carl Zeiss Microimaging, Inc., Thornwood, NY, USA). A line tool (AxioVision LE 29A software version 4.6.3.) measures horizontal ABR between the cementoenamel junction and alveolar bone crest [10,39,40]. Three examiners were blinded to measure the ABR of polybacterial and sham infection groups.

### RNA isolation and purification

Mandible from each mouse was homogenized with handheld rotor-stator homogenizer and sterile individual TissueRuptor disposable probes (Qiagen; Germantown, MD, USA) in the presence of lysis/binding buffer from mirVana miRNA isolation kit (Thermo Fisher Scientific, Waltham, MA, USA; Catalog number: AM1560). Samples were processed following the manufacturer’s protocol, and total RNA was extracted. The concentration and purity of the isolated total RNA was determined at 230, 260, and 280 nm in the NanoDrop Spectrophotometer (ND-1000 Thermo Fisher Scientific). Total RNA was used for RT-qPCR experiments [34].

### Synthesis of cDNA

The purified total RNA was reverse transcribed (20 ng in each sample) into cDNA using miRNA-specific primers and reverse transcription (RT) master mix from TaqMan microRNA reverse transcription kit (Thermo Fisher Scientific, USA). The synthesized cDNA is stored at –20°C for further qPCR experiments. The primer sequences for the 15 different miRNAs are shown in Table 2.

**TABLE 2.**
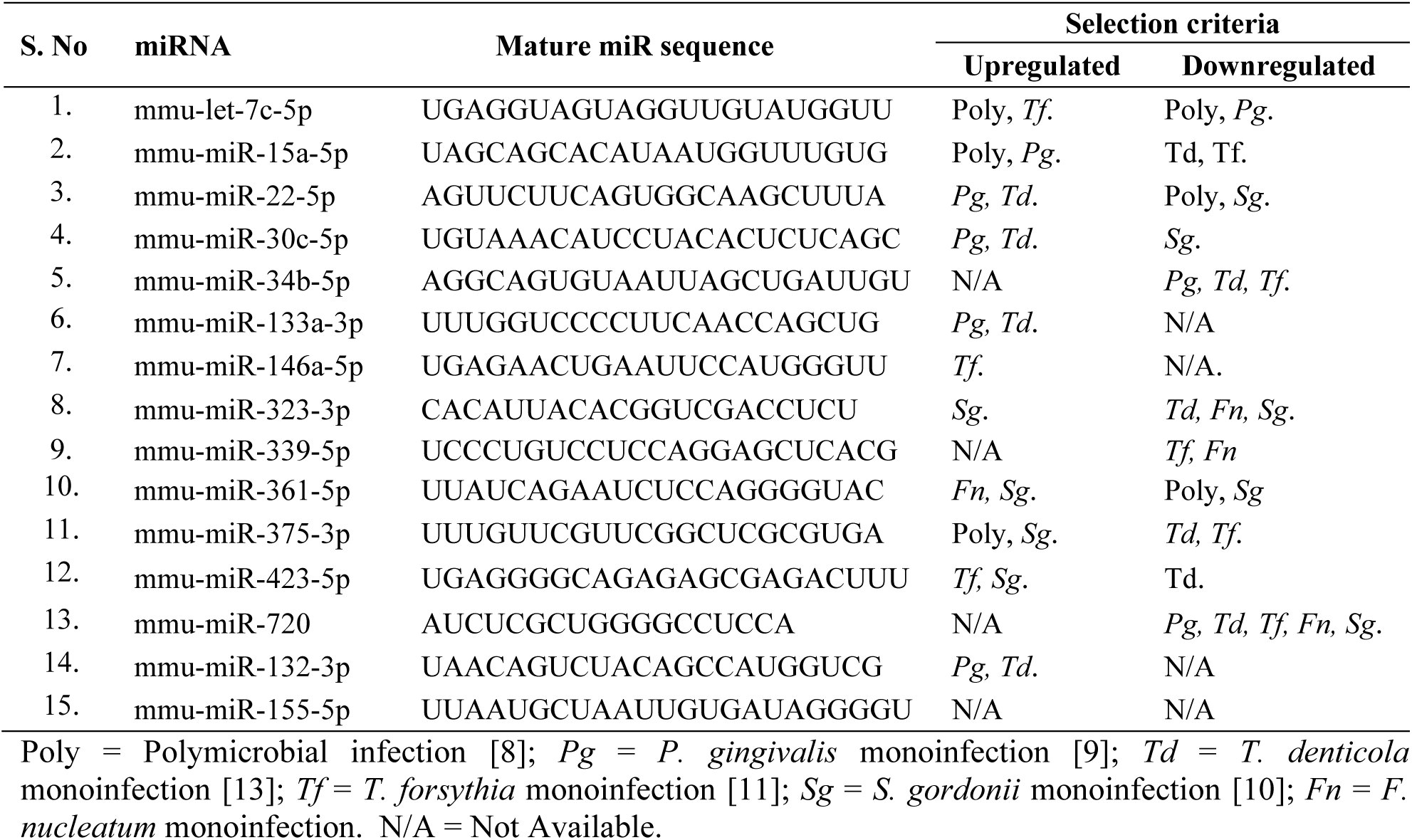
Primer sequences of the 15 selected candidate miRNAs.

### RT-qPCR assay

The reverse transcribed cDNA (2µl of synthesized cDNA per reaction) was used for qPCR experiments. Reagents used for the qPCR assay were purchased from Life Technologies (Carlsbad, CA). qPCR reactions were performed using the StepOne Real-Time PCR System (Applied Biosystems). Samples run in duplicates. The protocol includes an initial denaturation step at 95°C for 1 min, followed by 40 cycles consisting of denaturation at 95°C for 10 sec denaturation, annealing at 50°C for 20 sec, and extension at 72°C for 25 sec. snoRNA202 was used as an endogenous control for miRNA expression analysis. miRNA levels were normalized to snoRNA202 expression in mouse samples, and cycle threshold (Ct) values, corresponding to the PCR cycle number at which fluorescence emission reaches a threshold above baseline emission exceeds a baseline threshold, were determined relative miRNA expression was calculated using the 2**^−ΔΔCt^** method [41,42].

### Receiver operating characteristic (ROC) curve analysis

The miRNAs associated with periodontitis (diagnostic miRNAs) were evaluated using the receiver operating characteristics curve. RT-qPCR data was analyzed by using the MedCalc® Statistical Software version 22.026 (MedCalc Software Ltd, Ostend, Belgium; https://www.medcalc.org; 2024), ROC curve analysis was done, and optimal cutoff values for periodontal miRNAs was determined. The calculated sensitivity and specificity were recorded [43].

### Statistical analysis

The statistical significance of ABR data was determined using One-way ANOVA with Dunnett’s multiple comparisons set, and analysis was performed using the statistical software Prism 9.4.1 (GraphPad Software, San Diego, CA, USA). The Mann–Whitney *U* test was used for IgG antibody. For the RT-qPCR data, statistical analysis was performed using the Mann-Whitney *U* test to determine significant differences in miRNA expression between polymicrobial infection-treated and sham-infection mice. Data are presented as the mean ± standard deviation (SD), and *p* < 0.01 to *p* < 0.05 was considered statistically significant.

## RESULTS

### Gingival colonization of bacteria and host immune response

Analysis of gingival plaque swabs from each mouse (wild-type, TLR2^−/−^, and TLR4^−/−^) after bacterial infection showed the presence of bacterial-specific gene amplicons (16s rRNA) in agarose gel electrophoresis. Mice infected with early ecological colonizer *S. gordonii* showed >50% colonization for *S. gordonii* in the first infection cycle (1 IC) and 100% after the second infection cycle (2 IC), indicating that all mice were colonized with Gram+ *S. gordonii*. Gingival swabs from the *F. nucleatum* infection showed >50% colonization after the first IC and reached 100% colonization in the second IC. This confirmed the successful colonization of intermediate periodontal colonizers. Subsequently, 3 weeks after the infection with *P. gingivalis*, *T. denticola,* and *T. forsythia* (2 ICs), mice gingival swabs showed 30-60 % bacterial colonization for all three bacteria and reached 75-90% colonization after mice received 4^th^ infection cycle. None of the sham-infected mice were positive at any point for 16s rRNA of any of the five bacteria we analyzed. These results confirmed the successful colonization of five bacteria in the infected mice’s oral cavity (Table 3). Serum from the wild-type, TLR2^−/−^, and TLR4^−/−^ mice were evaluated for humoral/IgG immune response against the formalin-killed whole-cell antigens of *F. nucleatum, P. gingivalis, T. denticola,* and *T. forsythia*. A significant increase of Ig immune response was observed in infected mice groups; specifically the wild-type mice exhibited highly significant serum IgG response >1000 folds to *P. gingivalis* and *T. forsythia*, TLR2^−/−^ and TLR4^−/−^ mice showed significant IgG response to *F. nucleatum* and *P. gingivalis* (>90 folds), and TLR4^−/−^ mice had a robust level of IgG response to *P. gingivalis* (>700 folds) (Figure 1D1-3).

**TABLE 3.** Distribution of gingival plaque samples positive for periodontal pathogens gDNA.

| Group/Mice/Infection | Positive gingival plaque samples (n =16 mice) |  |  |  |  |  |
| --- | --- | --- | --- | --- | --- | --- |
|  | 2 Weeks<br>(Sg) | 4 Weeks<br>(Sg) | 6 weeks<br>(Fn) | 8 weeks<br>(Fn) | 10 weeks<br>(Pg/Td/Tf) | 14 weeks<br>(Pg/Td/Tf) |
| Gr I Wild-type Infection | 11 | 16 | 12 | 16 | 10/9/7 | 15/13/14 |
| Gr II Wild-type Sham-Infection | 0 | NC | 0 | NC | 0/0/0 | NC |
| Gr III TLR2<sup>-/-</sup> Infection | 13 | 16 | 11 | 16 | 8/6/9 | 14/12/13 |
| Gr IV TLR2<sup>-/-</sup> Sham-Infection | 0 | NC | 0 | NC | 0/0/0 | NC |
| Gr V TLR4<sup>-/-</sup> Infection | 10 | 16 | 9 | 16 | 10/7/8 | 14/13/15 |
| Gr VI TLR4<sup>-/-</sup> Sham-Infection | 0 | NC | 0 | NC | 0/0/0 | NC |

NC- not collected gingival plaque samples. Sg – *S. gordonii*, Fn-*F. nucleatum*, Pg – *P. gingivalis*, Td- *T. denticola*, Tf- *T. forsythia*.

### Deficiency of TLR2 and TLR4 signaling reduces Alveolar Bone Resorption (ABR)

Polymicrobial infection in wild-type mice resulted in a significant increase in ABR on the mandible lingual (*p* < 0.0001) and maxilla buccal side (*p* < 0.01) than sham-infected mice (Figure 1B and C1). In contrast, the polymicrobial infection did not induce bone resorption in alveolar bone in the mandible lingual, maxilla palatal, and maxilla buccal sides in TLR2^−/−^ mice and ABR was not significantly higher in the infected animals than in sham-infected mice (Figure 1B and C2). Similarly, polymicrobial infection induced no ABR in the mandible lingual, maxilla palatal, and maxilla buccal side in TLR4^−/−^ mice, and ABR was not significantly higher in infected mice than the sham-infected mice (Figure. 1B & C3). These findings suggested that resorption of alveolar bone is closely related to gingival inflammation, and mice lacking TLR2 and TLR4 receptors exhibit dampened periodontal inflammatory responses.

### Deficiency of TLR2 and TLR4 signaling reduces gingival inflammation

Histological examination of the mouse mandible in the infected TLR2^−/−^ and TLR4^−/−^ mice demonstrated minimal apical migration of junctional epithelium (JE), gingival hyperplasia, and mild inflammatory cellular infiltration in connective tissue (Figure 1E2, E3) when compared to sham-infected TLR2^−/−^ and TLR4^−/−^ mice as well as both male and female mice.

These results suggest that TLR2 and TLR4 signaling are critical and play an important role in inflammatory response against polybacterial infection. In contrast, histological examination of mouse mandible in the infected wild-type mice showed moderate apical migration of JE, gingival hyperplasia, and inflammatory cellular infiltration in the connective tissue (Figure 1E1) when compared to sham-infected mice as well as both male and female mice.

### Polymicrobial infection-induced systemic translocation of bacteria to distal organs

To examine whether periodontal bacteria *S. gordonii*, *F. nucleatum, P. gingivalis, T. denticola,* and *T. forsythia* colonized/infected the gingival margins of molar teeth translocated intravascularly to multiple internal organs, we isolated bacterial genomic DNA from the heart, aorta, brain, liver, kidney, and lung and PCR was performed to detect the presence of specific bacterial gDNA. Bacteria-specific genomic DNA from all five oral microbes (*Sg/Fn/Pg/Td/Tf*) were identified in the heart and lungs, and *P. gingivalis, T. denticola* in the brain in wild-type mice (Table 4). Whereas *P. gingivalis* and *T. forsythia* gDNA were observed in all organs except the spleen in TLR2^−/−^ mice. The gDNA of *S. gordonii* was observed in the infected mice’s lungs and kidneys in TLR2^−/−^ mice. The gDNA of *F. nucleatum* was detected in the heart, *P. gingivalis* in the lungs, and *T. forsythia* in the kidney of infected TLR4^−/−^ mice (Table 4).

**TABLE 4.** PCR test results for bacterial dissemination to distal organs.

| Group/Mice | PCR Positive samples (n=16) |  |  |  |  |  |
| --- | --- | --- | --- | --- | --- | --- |
|  | Heart<br>*Sg/Fn/Pg/Td/Tf* | Lungs<br>*Sg/Fn/Pg/Td/Tf* | Brain<br>*Sg/Fn/Pg/Td/Tf* | Liver<br>*Sg/Fn/Pg/Td/Tf* | Kidney<br>*Sg/Fn/Pg/Td/Tf* | Spleen<br>*Sg/Fn/Pg/Td/Tf* |
| I Wild-type | 13/7/9/11/10 | 8/9/7/2/11 | 0/0/5/3/0 | 0/0/0/0/3 | 8/2/0/0/5 | 2/0/0/0/1 |
| II Sham | 0/0/0/0/0 | 0/0/0/0/0 | 0/0/0/0/0 | 0/0/0/0/0 | 0/0/0/0/0 | 0/0/0/0/0 |
| III TLR2<sup>-/-</sup> | 0/0/7/3/12 | 12/0/0/0/16 | 0/0/8/0/8 | 0/0/8/0/7 | 3/0/1/0/10 | 0/0/0/0/0 |
| IV Sham | 0/0/0/0/0 | 0/0/0/0/0 | 0/0/0/0/0 | 0/0/0/0/0 | 0/0/0/0/0 | 0/0/0/0/0 |
| V TLR4<sup>-/-</sup> | 0/6/0/0/0 | 0/0/5/0/0 | 0/0/0/0/0 | 0/0/0/0/0 | 0/0/0/0/2 | 0/0/0/0/0 |
| VI Sham | 0/0/0/0/0 | 0/0/0/0/0 | 0/0/0/0/0 | 0/0/0/0/0 | 0/0/0/0/0 | 0/0/0/0/0 |

Group I = Wild-type mice / ETSPPI (Ecological Time-sequential Polymicrobial Periodontal Infection). Group II = Wild-type mice / Sham-Infection. Group III = TLR2<sup>-/-</sup> mice / ETSPPI Infection. Group IV = TLR2<sup>-/-</sup> mice / Sham-Infection. Group V = TLR4<sup>-/-</sup> mice / ETSPPI Infection. Group VI = TLR4<sup>-/-</sup> mice / Sham-Infection.

**TABLE 5.** Polymicrobial infection-induced upregulated miRNAs reported functions and target genes.

| miRNA | P value | AUC value | Target Functions | Target genes |
| --- | --- | --- | --- | --- |
| **Upregulated miRNAs in the wild-type mice** |  |  |  |  |
| **miR-146a-5p** | 0.01 | 0.97 | Upregulated in the gingiva of periodontitis patients [44-46] and saliva of periodontitis patients [47]. Upregulated in the MM6 cells differentiated in the presence of LPS (TLR4) or zymosan (TLR2) for 96 h [48]. | 29<br>(*PLN*,<br>*Relb*,<br>*Cd93*,<br>*Sgk3*,<br>*Irak2*) |
| **miR-15a-5p** | 0.05 | 0.87 | Upregulated in gingival crevicular fluid from periodontitis patients [49] and in serum of myocardial fibrosis patients, serum, and myocardial tissues of Ang-II-treated mice [50]. **Promoting cancer cell invasion and metastasis capabilities through miR-15a-5p/CXCL17 axis** [51]. Significantly increased in the exosomes from patients with advanced carotid atherosclerosis [52]. Upregulated in the CSF of Alzheimer's disease [53]. | 519<br>(*Tacc1*,<br>*Bcl2l1*,<br>*Scn4b*,<br>*Atg14*,<br>*Fkrp*) |
| **TLR2<sup>-/-</sup> mice** |  |  |  |  |
| **miR-146a-5p** | 0.01 | 0.95 | Described in the wild-type mice. |  |
| **miR-15a-5p** | 0.05 | 0.82 | Described in the wild-type mice. |  |
| **TLR4<sup>-/-</sup> mice** |  |  |  |  |
| miR-22-5p | 0.05 | 0.8 | Upregulated in the plasma samples of acute myocardial infarction patients [54,55]. | - |
| miR-30c-5p | 0.01 | 0.88 | **Upregulated in gingival crevicular fluid from chronic periodontitis patients** [44]. Reported in chronic heart failure [56]. Upregulated in spared nerve injury rat prefrontal cortex [57]. Downregulated in atherosclerosis [58]. Potential biomarker in the acute rejection after heart transplantation [59]. Downregulated in CSF and sera of Cryptococcosis patients [60]. Upregulated in the urine of post-cardiac-surgery patients [61]. | 47<br>(*Egfr*,<br>*Twf1*,<br>*Pgr*,<br>*Six4*,<br>*Ing1*,<br>*Reep4*) |
| **miR-146a-5p** | 0.01 | 0.94 | Described in the wild-type mice. |  |
| miR-323-5p | 0.05 | 0.85 | **Upregulated in obese periodontitis patients** [62]. Positive regulator of myogenesis [63], a biomarker for cardiomyopathy [64]. Upregulated in blood samples of coronary heart disease patients and elevated in coronary arteries of coronary heart disease rat model [65]. | 25<br>(*Pira1*,<br>*Pira2*,<br>*Tyw1*,<br>*Atf7*,<br>*Iars2*) |
| miR-361-5p | 0.01 | 0.96 | Upregulated in acute coronary syndrome [66]. Downregulated in *F. nucleatum*-infected colorectal cancer cell lines (HT-29, HCT 116) [67]. Upregulated in the serum of Acute myocardial injury patients [68]. Upregulated in the serum of the *M. tuberculosis* patients [69]. Reported as a biomarker for clinical diagnosis of atherosclerosis [70]. | 15<br>(*Ctbp2*,<br>*Tfam*,<br>*Gid4*,<br>*Ssr1*,<br>*Eea1*) |
| miR-375-3p | 0.01 | 0.95 | Upregulated in gingival crevicular fluid from chronic periodontitis patients [44], *S. gordonii* induced periodontitis [10]. Upregulated in mice exposed to ionizing radiation [71]. Upregulated in hearts of the transverse aortic constriction rat model [72]. Negative regulator of | 19<br>(*Mtpn*,<br>*Akap2*,<br>*Atrn*, |
|  |  |  | osteogenesis [73]. Decreased expression was reported in the MPTP-mediated PD mouse model [74]. | *Rpsa*,<br>*Yap1* |
| miR-720 | 0.01 | 0.98 | The most expressed miRNA in chronic and aggressive periodontitis patients [75]. Upregulated in HBV-specific CD8+ T cells [76]. Upregulated in the serum of hepatitis patients [77]. Downregulated in cardiac function for patients after surgery [78]. | 9 (*Pon2*,<br>*Hlcs*,<br>*Agap1*,<br>*Ppp6r1*) |
| let-7c-5p | 0.01 | 0.881 | Upregulated in the Bacillus Calmette Guérin (BCG) infected macrophage cell lines [79]. Inhibiting the microglial activation in the cerebral ischemia condition [80]. | 21 (*Nf2*,<br>*Cts8*,<br>*Lins*,<br>*Msi2*,<br>*Myc*) |

### miR-146a-5p, miR-15a-5p, miR-132-3p are elevated and Let-7c, miR-22 are decreased in wild-type mice

Fifteen miRNA expressions in the mandibles from polymicrobial-infected wild-type and sham-infected mice were analyzed using the RT-qPCR. Three miRNAs miR-132-3p (p < 0.0001; Figure 2A), miR-146a-5p (p ≤ 0.01; Figure 2B), and miR-15a-5p (p ≤ 0.05; Figure 2C) were upregulated with a ROC -area under the curve for miR-132-3p (AUC: 1.00), miR-146a-5p (AUC: 0.97) and miR-15a-5p (AUC: 0.87) in the polymicrobial infected wild-type mice compared to sham-infected mice. miR-132-3p has the highest AUC value, followed by miR-146a-5p and miR-15a-5p. The miRNAs of miR-let-7c-5p (Figure 2D) and miR-22-5p (Figure 2E) were downregulated. The remaining ten miRNA expressions showed no difference between infection and sham infection (Supplementary Figure S1).

**FIGURE 2.**
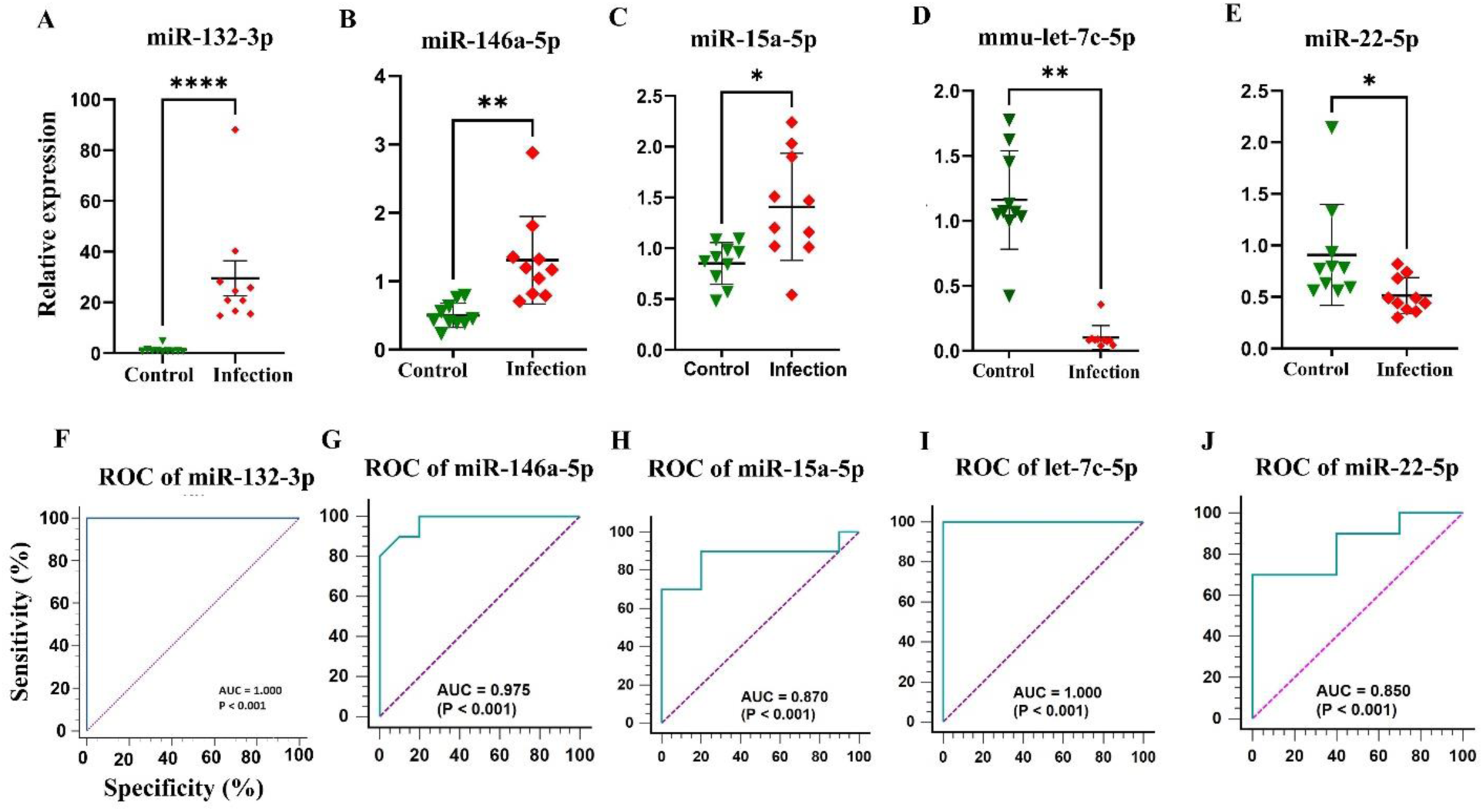
Relative expression levels were determined by quantitative reverse transcription polymerase chain reaction for selected microRNAs (miRNAs) in the mandibles of the polymicrobial-infected wild-type and sham-infected mice. ROC curve of miRNAs that correlates with polybacterial infection-induced periodontitis. (E) mmu-let-7c-5p ROC curve with *p* < 0.001. (F) miR-15a-5p ROC curve with *p* < 0.001. (G) miR-22-5p ROC curve with *p* < 0.001. (H) miR-146a-5p ROC curve with *p* < 0.001.

**FIGURE 3.**
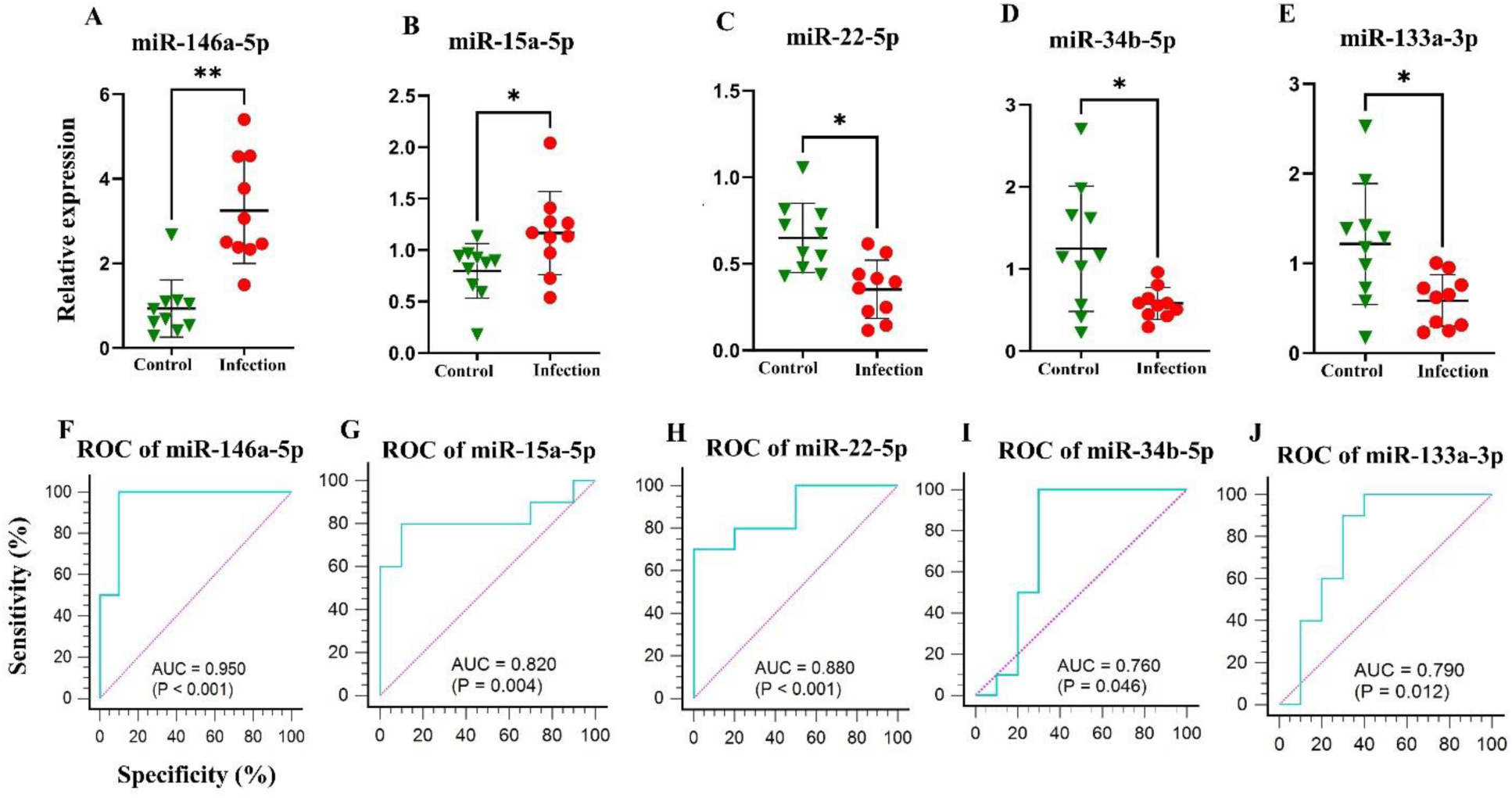
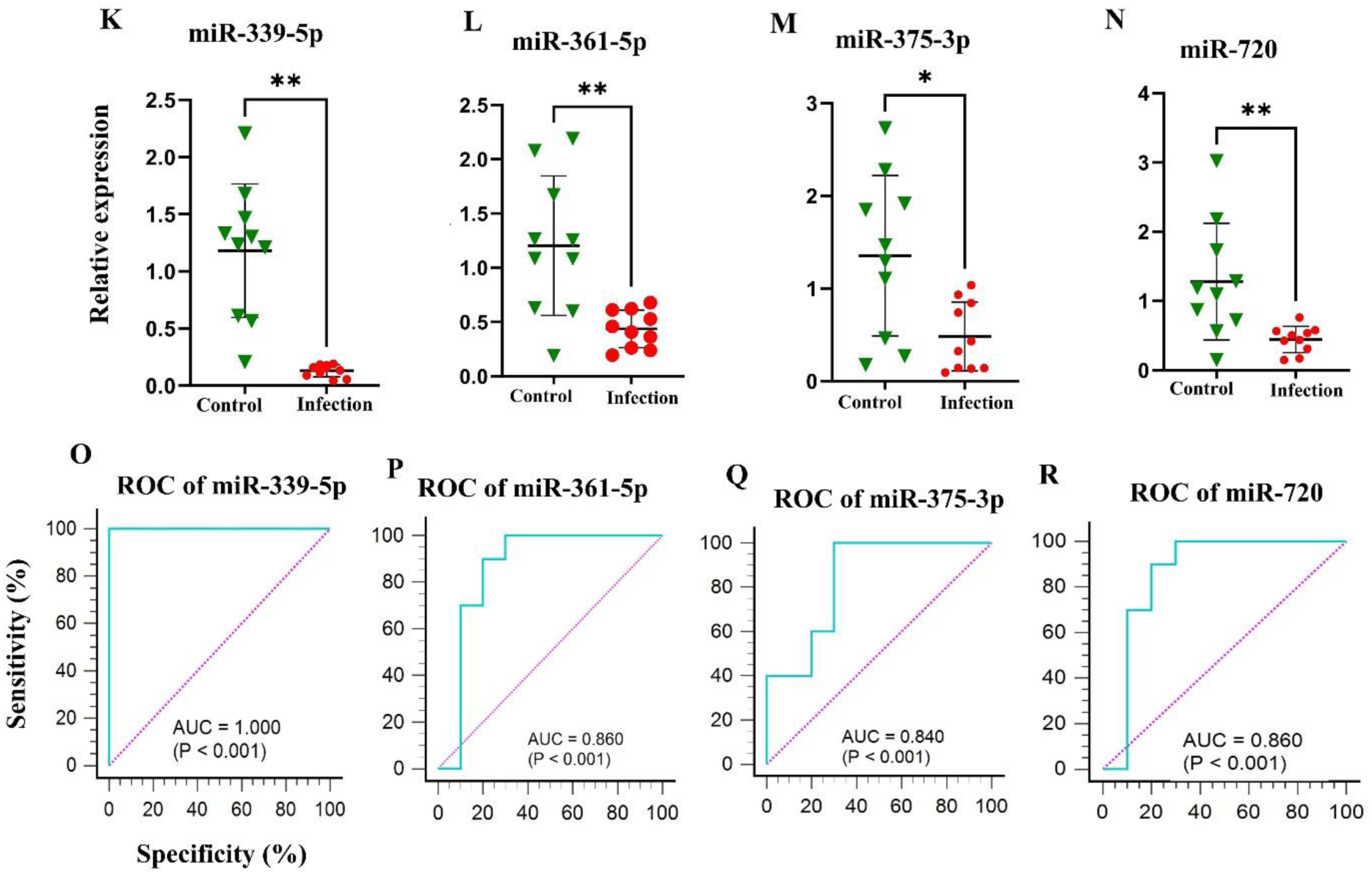
Relative expression levels were determined by quantitative reverse transcription polymerase chain reaction for selected microRNAs (miRNAs) in the mandibles of the polymicrobial infected TLR2^−/−^ mice vs TLR2^−/−^ sham infected mice. ROC curve of miRNAs that correlates with polybacterial infection-induced periodontitis.

### miR-146a-5p, miR-15a are elevated and 7 miRNAs downregulated in TLR2^−/−^ mice

The expression profile of 15 miRNAs analyzed in the mandibles of TLR2^−/−^ mice was distinct from the wild-type mice. Notably, miR-146a-5p (p ≤ 0.01; Figure 2A) and miR-15a-5p (p ≤ 0.05; Figure 2B) were upregulated with an area under the curve for miR-146a-5p (AUC: 0.95) and miR-15a-5p (AUC: 0.82). While both wild-type and TLR2^−/−^ mice showed similar upregulation of these two miRNAs, their TLR2^−/−^ expression patterns were altered. miR-146a-5p has the highest AUC value compared to miR-15a-5p. The miRNAs of miR-22-5p, miR-34b-5p, miR-133a-3p, miR-339-5p, miR-361-5p, miR-375-3p and miR-720 was significantly downregulated in TLR2^−/−^ mice mandibles. The expression of the remaining miRNA showed no significant variation in infection and sham infection (Supplementary Figure S2).

### miR-146a-5p and other 7 miRNAs elevated in mandibles of TLR4^−/−^ mice

The elevated expression levels of miR-146a-5p (p ≤ 0.01; Figure 4A) with an area under the curve (AUC: 0.94) were observed in TLR4^−/−^ mice and were consistent with the infection groups in the wild-type mice and TLR2^−/−^ mice. Five of the miRNAs, miR-30c-5p, miR-361-5p, miR-375-3p, miR-720, and mmu-let-7c-5p were upregulated with a *p* ≤ 0.01 value, and two miRNAs miR-22-5p, and miR-323-3p, with a *p* ≤ 0.05 value. The AUC values for upregulated miRNAs includes; miR-30c-5p (Fig 4F; AUC: 0.88), miR-361-5p (Fig 4G; AUC: 0.96), miR-375-3p (Fig 4H; AUC: 0.95), miR-720 (Fig 4M; AUC: 0.98), mmu-let-7c-5p (Fig 4N; AUC: 0.881), miR-22-5p (Fig 4O; AUC: 0.8), and miR-323-3p (Fig 4P; AUC: 0.85). The remaining seven miRNA expressions showed no difference between infection and sham infection (Supplementary Figure S3).

**Figure 4.**
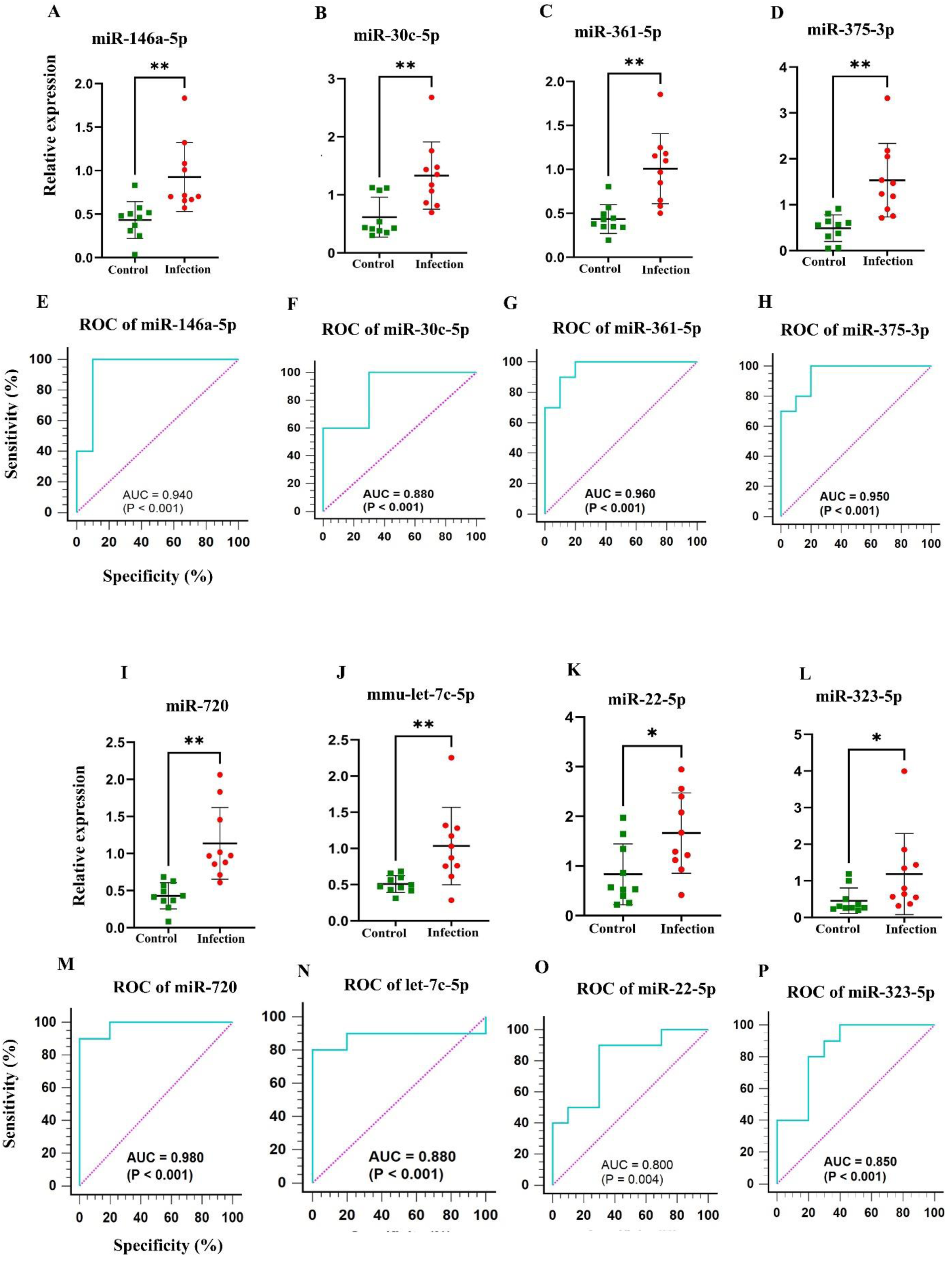
Relative expression levels were determined by quantitative reverse transcription polymerase chain reaction for selected microRNAs (miRNAs) in the mandibles of the polymicrobial infected TLR4^−/−^ mice vs TLR4^−/−^ sham infected mice. ROC curve of miRNAs that correlates with polybacterial infection-induced periodontitis.

### Differentially Expressed miRNAs and Functional Pathways Analysis

Predicted functional pathway analysis of the DE miRNAs using KEGG in the polymicrobial infected wild-type mice mandibles identified several pathways, such as Wnt-signaling pathway, focal adhesion pathway, VEGF signaling pathway, B- and T-cell receptor signaling pathway, Axon guidance, TGF-beta signaling pathway, HTLV-1 infection, PI3K-Akt pathway, MAPK signaling pathway, TLR signaling pathway, and osteoclast differentiation pathway (Figure 5A). Three upregulated miRNAs of wild-type mice have altered the expression of ten genes in the TLR signaling pathway (Figure 6A). The two DE upregulated miRNAs from TLR2^−/−^ infected mice mandibles associated with several pathways such as pluripotency of stem cells, TGF-beta signaling pathway, PI3K-Akt pathway, TLR signaling pathway, and different pathways of cancers (Figure 5B). The two upregulated miRNAs in TLR2^−/−^ mice altered nine genes’ expression in the TLR signaling pathway (Figure 6B). Seven downregulated miRNAs of the TLR2 mice have altered 14 gene expressions in the bacterial invasion of the epithelial cell pathway (Figure 5C). Bacterial invasion of epithelial cells pathway shows the altered gene expression depicted in (Figure 6C). The eight DE upregulated miRNAs from the TLR4^−/−^ infected mice mandibles are associated with the pathways of focal adhesion, MAPK signaling pathway, regulating pluripotent stem cells pathway, platelet activation pathway, TGF-beta signaling pathway, osteoclast differentiation pathway, and ubiquitin-mediated proteolysis pathway (Figure 5D). The upregulated miRNAs from the TLR4^−/−^ infected mice have altered 27 genes in the osteoclast differentiation pathway.

**FIGURE 5A.**
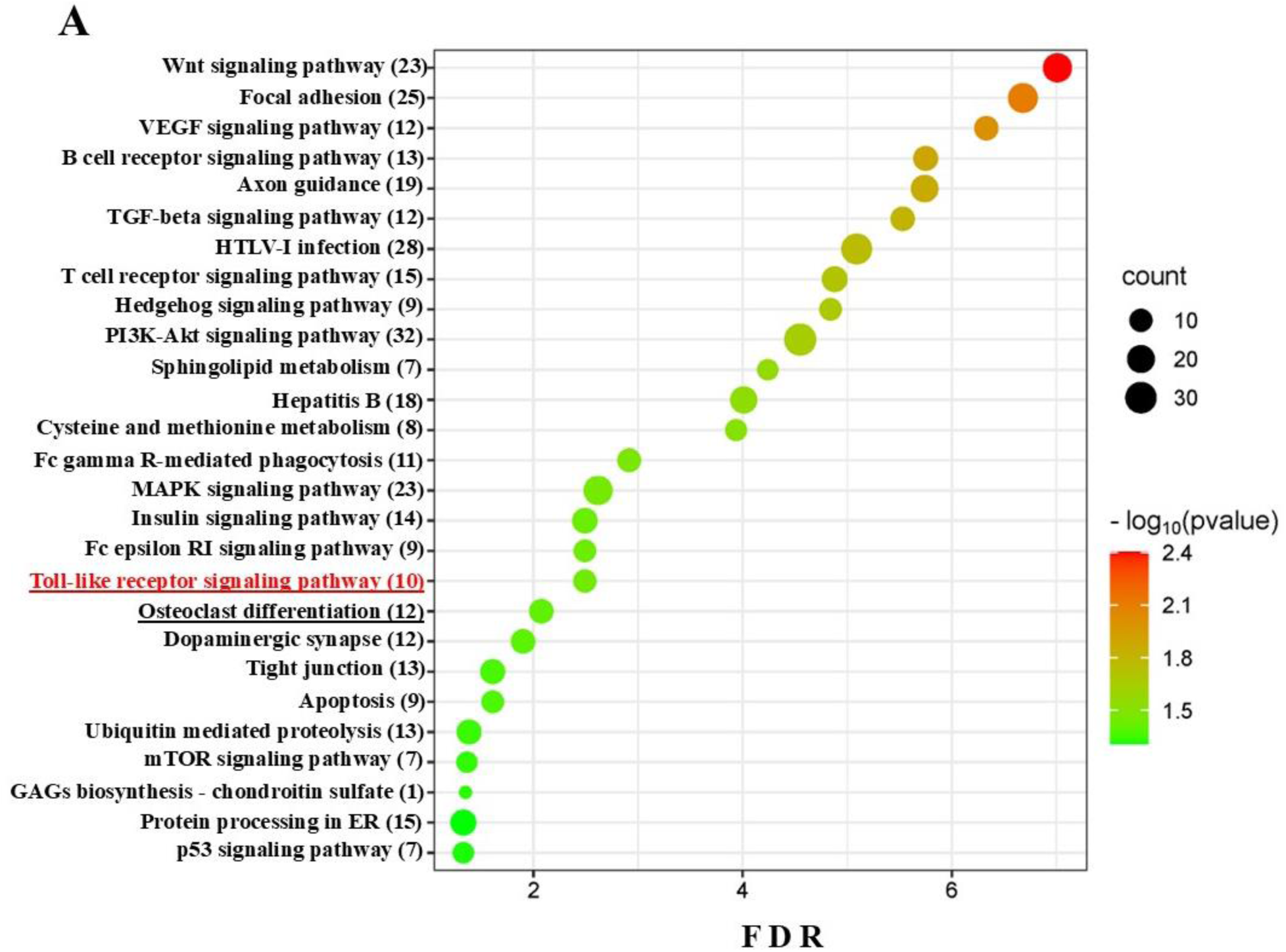
Bubble plot for upregulated miRNAs in wild-type mice. The predicted functional pathway analysis of DE miRNAs from polymicrobial infected wild-type mice mandibles. Bubble plot of KEGG analysis on predicted target genes of DE miRNAs in polymicrobial infected mice compared to sham-infected mice. The KEGG pathways are displayed on the y-axis showing the number of genes altered in the pathway in brackets, and the x-axis represents the false discovery rate (FDR), which means the probability of false positives in all tests. The size and color of the dots represent the number of predicted genes and corresponding p-value, respectively. Ten genes were shown to be altered in the Toll-like receptor signaling pathway.

**FIGURE 5B.**
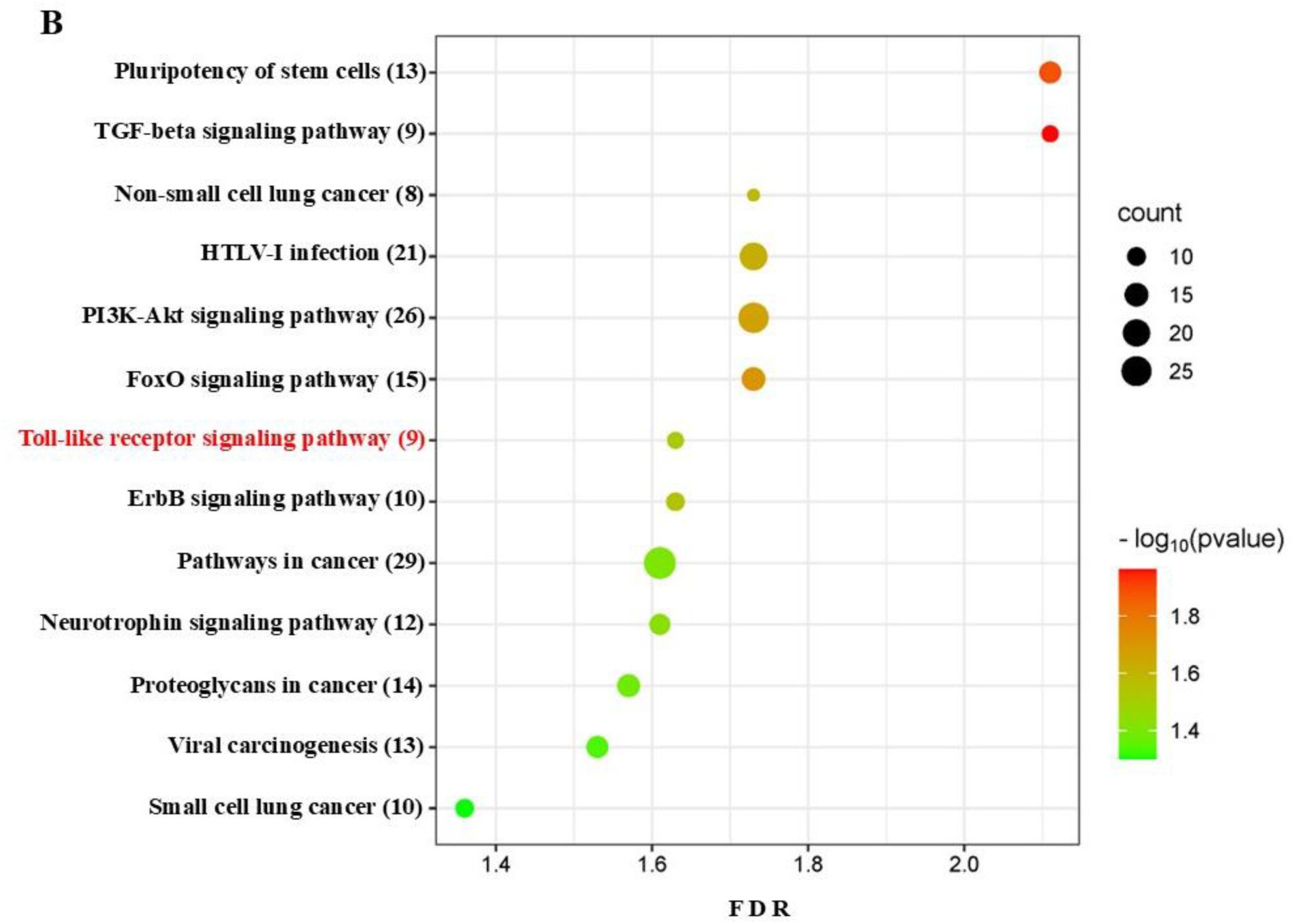
Bubble plot for upregulated miRNAs in wild-type and TLR2^−/−^ mice. The predicted functional pathway analysis of DE miRNAs from polymicrobial infected wild-type and TLR2^−/−^ mice mandibles. Bubble plot of KEGG analysis on predicted target genes of DE miRNAs in polymicrobial infected mice compared to sham-infected mice. The KEGG pathways are displayed on the y-axis, showing the number of genes altered in the pathway in brackets, and the x-axis represents the false discovery rate (FDR), which means the probability of false positives in all tests. The size and color of the dots represent the number of predicted genes and corresponding p-value, respectively. Nine genes were shown to be altered in the Toll-like receptor signaling pathway.

**Figure 5C.**
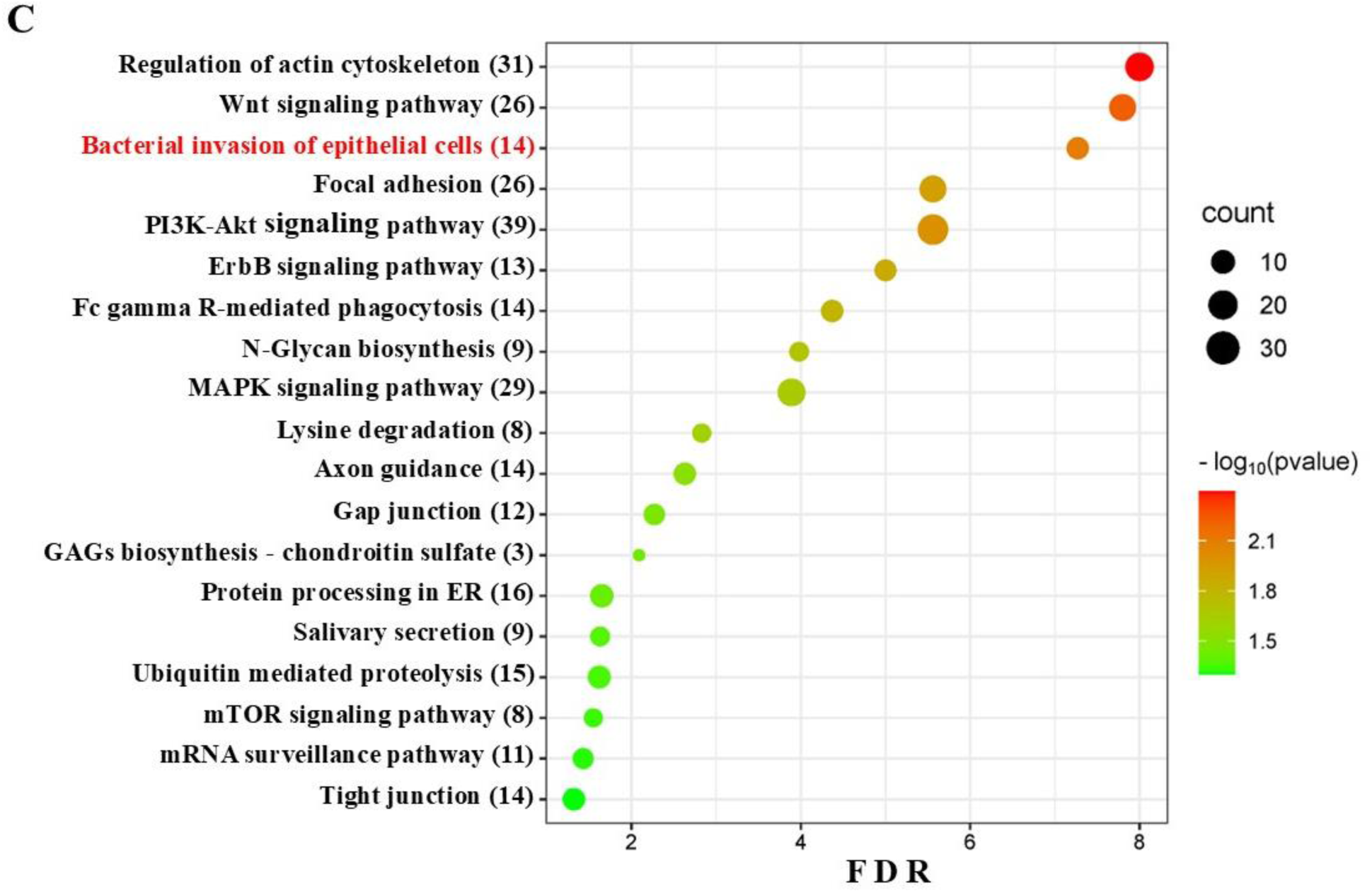
Bubble plot for downregulated miRNAs in TLR2^−/−^ mice. The predicted functional pathway analysis of DE miRNAs from polymicrobial infected TLR2^−/−^ mouse mandibles. Bubble plot of KEGG analysis on predicted target genes of DE miRNAs in polymicrobial infected mice compared to sham-infected mice. The KEGG pathways are displayed on the y-axis, showing the number of genes altered in the pathway in brackets, and the x-axis represents the false discovery rate (FDR), which means the probability of false positives in all tests. The size and color of the dots represent the number of predicted genes and corresponding p-value, respectively.

**FIGURE 5D.**
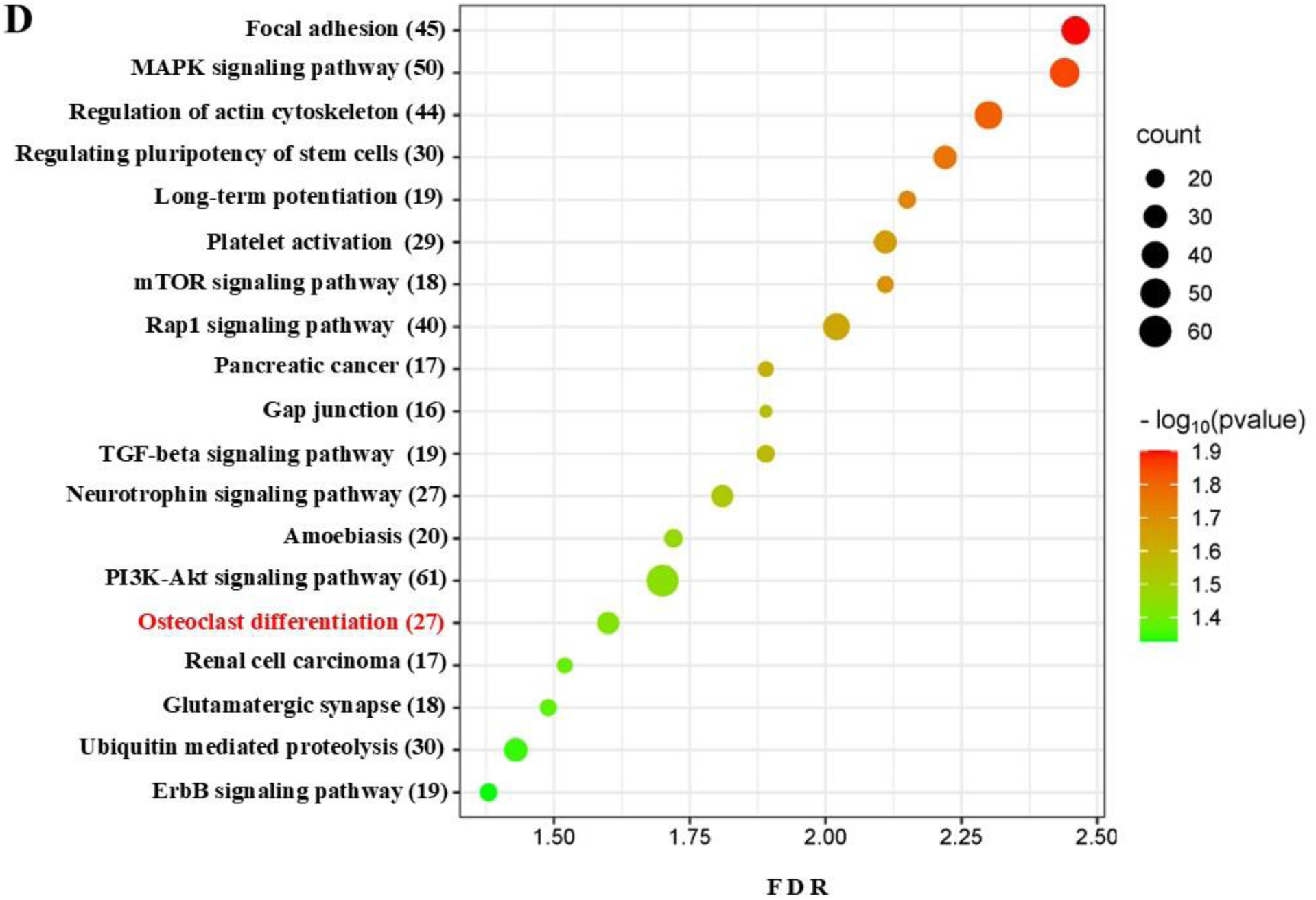
Bubble plot for upregulated miRNAs in TLR4^−/−^ mice. The predicted functional pathway analysis of DE miRNAs from polymicrobial infected TLR4^−/−^ mouse mandibles. Bubble plot of KEGG analysis on predicted target genes of DE miRNAs in polymicrobial infected mice compared to sham-infected mice. The KEGG pathways are displayed on the y-axis, showing the number of genes altered in the pathway in brackets, and the x-axis represents the false discovery rate (FDR), which means the probability of false positives in all tests. The size and color of the dots represent the number of predicted genes and corresponding p-value, respectively. 27 genes are known to be altered in the Osteoclast differentiation pathway.

**FIGURE 6A.**
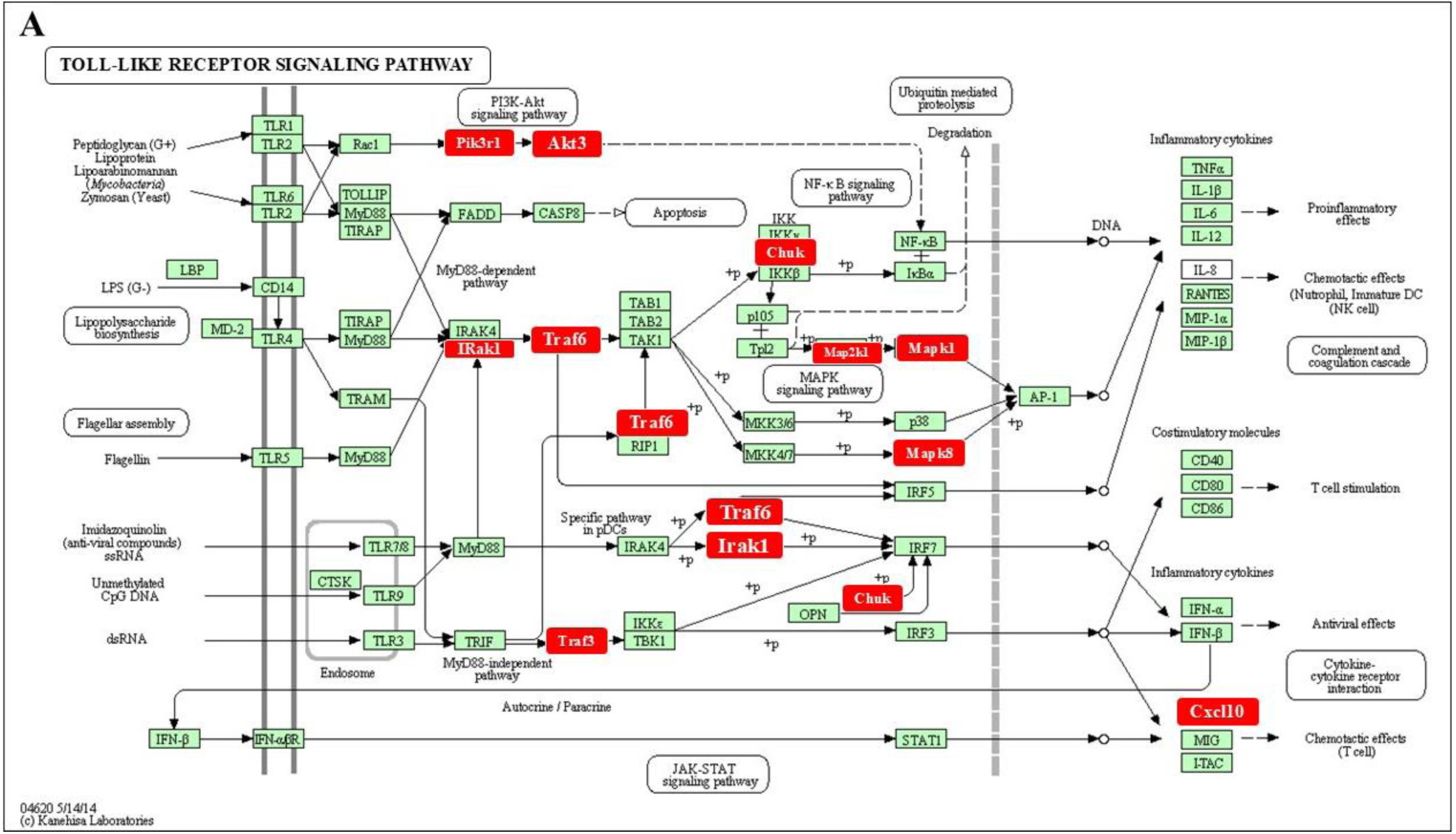
Upregulated miRNAs in wild-type mice altering 10 genes in Toll-like receptor signaling pathway. Significantly differentially expressed genes (identified by KEGG) participate in the TLR signaling pathway. Red boxes indicate significantly increased expression based on miRNA profiles from NanoString analysis. Green boxes indicate no change in gene expression. Specific families of pattern recognition receptors are responsible for detecting microbial pathogens and generating innate immune responses. Toll-like receptors (TLRs) are membrane-bound receptors identified as homologs of Toll in Drosophila. Mammalian TLRs are expressed on innate immune cells, such as macrophages and dendritic cells, and respond to Gram-positive or Gram-negative bacteria membrane components. Pathogen recognition by TLRs provokes rapid activation of innate immunity by inducing the production of proinflammatory cytokines and upregulation of costimulatory molecules. TLR signaling pathways are separated into two groups: a MyD88-dependent pathway that leads to the production of proinflammatory cytokines with quick activation of NF-kB and MAPK and a MyD88-independent pathway associated with the induction of IFN-beta and IFN-inducible genes, and maturation of dendritic cells with slow activation of NF-kB and MAPK.

**FIGURE 6B.**
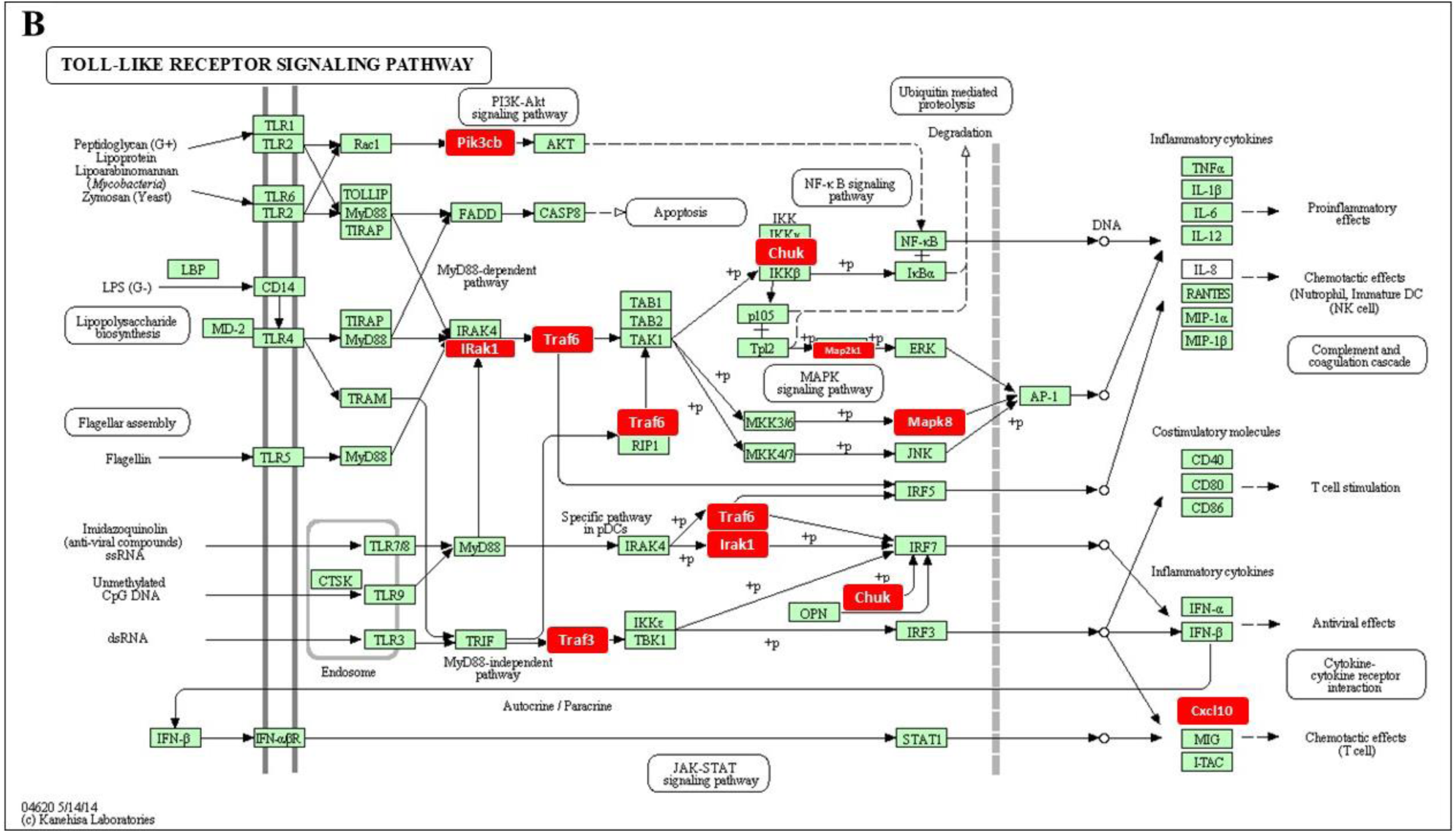
Upregulated miRNAs in TLR2^−/−^ mice altering 9 genes in TLR signaling pathway. Significantly differentially expressed genes (identified by KEGG) participate in the toll-like receptor signaling pathway. Red boxes indicate significantly increased expression based on miRNA profiles from NanoString analysis. Green boxes indicate no change in gene expression. Specific families of pattern recognition receptors are responsible for detecting microbial pathogens and generating innate immune responses. TLRs are membrane-bound receptors identified as homologs of Toll in Drosophila. Mammalian TLRs are expressed on innate immune cells, such as macrophages and dendritic cells, and respond to the membrane components of Gram-positive or Gram-negative bacteria. Pathogen recognition by TLRs provokes rapid activation of innate immunity by inducing the production of proinflammatory cytokines and upregulation of costimulatory molecules. TLR signaling pathways are separated into two groups: a MyD88-dependent pathway that leads to the production of proinflammatory cytokines with quick activation of NF-kB and MAPK and a MyD88-independent pathway associated with the induction of IFN-beta and IFN-inducible genes, and maturation of dendritic cells with slow activation of NF-kB and MAPK.

**FIGURE 6C.**
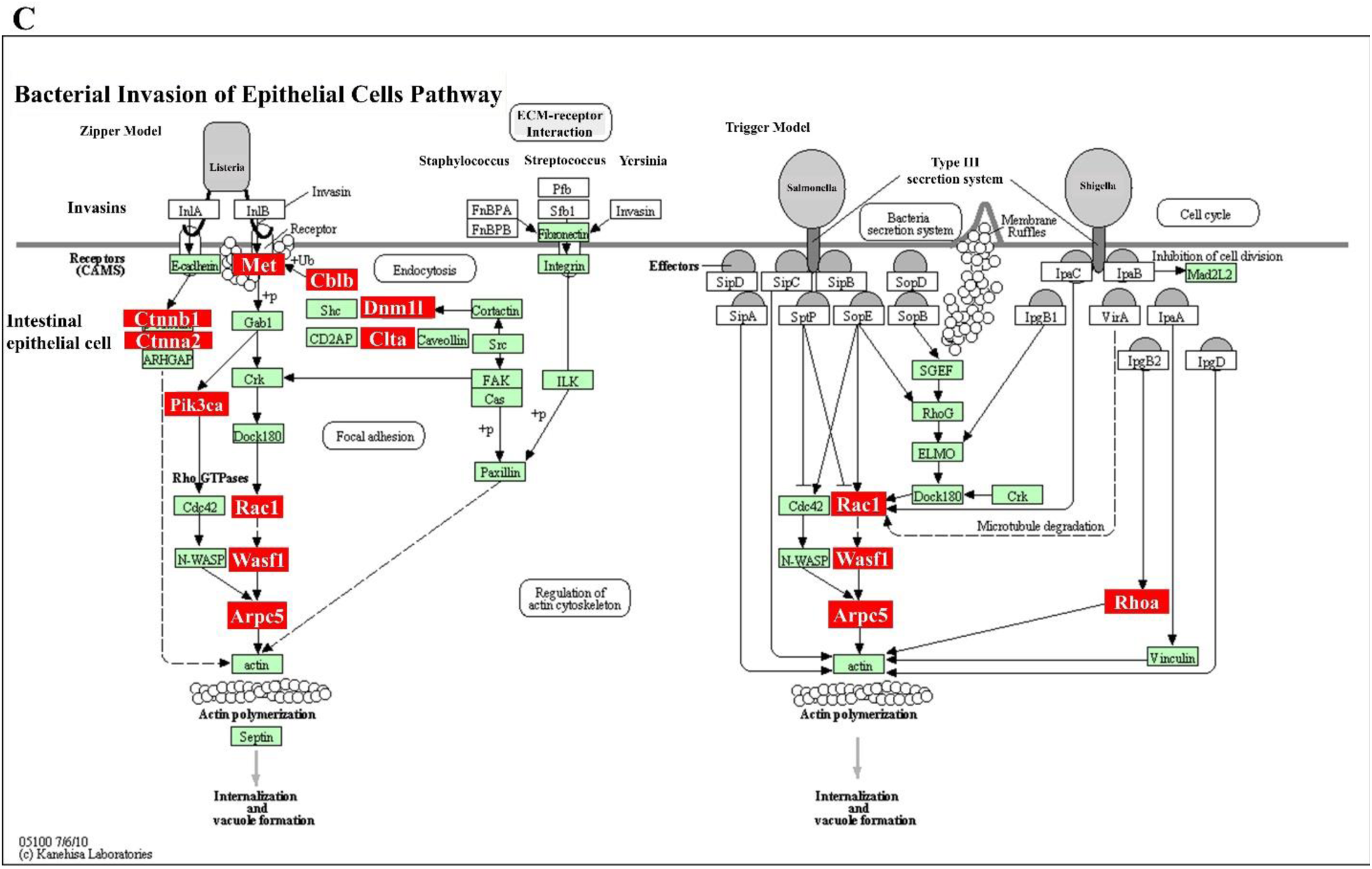
Downregulated miRNAs in TLR2^−/−^ mice altering 14 genes in bacterial invasion of epithelial cells pathway. Significantly, DE genes (identified by KEGG) participate in the TLR signaling pathway. Red boxes indicate significantly increased expression based on miRNA profiles from NanoString analysis. Green boxes indicate no change in gene expression. Many pathogenic bacteria can invade phagocytic and non-phagocytic cells and colonize them intracellularly, then become disseminated to other cells. Invasive bacteria induce their own uptake by non-phagocytic host cells (e.g. epithelial cells) using two mechanisms referred to as the zipper model and trigger model. *Listeria, Staphylococcus, Streptococcus*, and *Yersinia* are examples of bacteria that enter using the zipper model. These bacteria express proteins on their surfaces that interact with cellular receptors, initiating signaling cascades that result in close apposition of the cellular membrane around the entering bacteria. *Shigella* and *Salmonella* are examples of bacteria entering cells using the trigger model. These bacteria use type III secretion systems to inject protein effectors that interact with the actin cytoskeleton.

## DISCUSSION

The interplay between the dysbiotic microbial community and an abnormal host immune response, the gingival and periodontal tissue, contributes to the development of periodontitis [81]. The present study broadly investigated the colonization of five infected periodontal bacteria, infection-induced IgG antibody response, intravascular dissemination, histology, ABR, and miRNA analysis in the wild-type, TLR2^−/−^ and TLR4^−/−^ mice. Mice infected with ETSPPI had significant ABR in the wild-type mice but were not observed significantly among the TLR2^−/−^ and TLR4^−/−^ infected mice like our earlier study [22]. Five bacterial gDNA were detected in the heart, lungs, and limited for *P. gingivalis*, *T. denticola* in the brain in wild-type mice. Intravascular dissemination of five infected bacteria was very limited in TLR2^−/−^ and TLR4^−/−^ mice. Bacteria accessing blood circulation are commonly observed in the heart valves, liver, and spleen. Bacteria can also reach the brain hematogenously. Wild-type mice exhibited significant IgG responses for *P. gingivalis* and *T. forsythia* bacteria. while the infected TLR2^−/−^ and TLR4^−/−^ mice showed *P. gingivalis*-specific IgG antibody response like our earlier study Aravindraj et al, and Chukkapalli et al [8,22]. Histological examination of the mouse mandible in the infected TLR2^−/−^ and TLR4^−/−^ mice had minimal apical migration of junctional epithelium (JE), gingival hyperplasia, and mild inflammatory cellular infiltration in connective tissue.

Dysregulated or upregulated miRNAs identified as potential biomarkers for diagnosing the disease [82,83]. Several miRNAs are used as therapeutic molecules in disease conditions of cancer [84], type-2 diabetes mellitus [85], neurodegenerative diseases [86], and cardiovascular diseases [87] Diagnosing the early stages of PD will prevent the disease progression and improve the treatment.

We recently reported that periodontal bacteria can induce miRNA expression profiles, which differ from polymicrobial infections and mono-infection [8–11,13]. The selection of miRNA for RT-qPCR experiments depended on how uniquely miRNAs were expressed in 6 different infections [8–11,13]. For example, mmu-let-7c-5p was expressed in polymicrobial infection, *Tf*, and *Pg* mono-infection. miR-146a-5p was expressed in *Tf* mono-infection, miR-15a-5p was expressed in polymicrobial infection, and mono-infection of *Pg*, *Td*, and *Tf*. miR-132-5p was expressed in *Pg* and *Td* mono-infection. While miR-22-5p was expressed in polymicrobial infection and *Sg, Pg,* and *Td* mono-infection. miR-323-3p was expressed in the mono-infection of *Sg, Td, and Fn,* and miR-361-5p was expressed in polymicrobial infection and mono-infection of *Sg* and *Fn*. miR-375-3p was expressed in polymicrobial infection and mono-infection in *Td* and *Tf*. miR-720 was expressed in the mono-infection of *Sg, Fn, Pg, Td*, and *Tf* (Table 2). Therefore, in this study, we focused on the evaluation of 15 miRNAs such as let-7c-5p, miR-15a-5p, miR-22-5p, miR-30c-5p, miR-34b-5p, miR-133a-3p, miR-146a-5p, miR-323-3p, miR-339-5p, miR-375-3p, miR-361-5p, miR-423-5p, miR-720, miR-155-5p, and miR-132-3p revealed the expression difference between polymicrobial infection-treated mandibles and the healthy mice mandibles in the cohorts of wild-type, TLR2^−/−^, and TLR4^−/−^ mice. The miRNA analysis in this study provided PD-specific miRNA expression patterns in the wild-type mice and the miRNA hallmarks for periodontitis in the TLR-deficient mice.

This study found that miR-146a**-**5p, miR-15a-5p, and miR-132-3p were upregulated in wild-type infected mice. The miR-146a-5p, miR-15a-5p upregulated in TLR2^−/−^ infected mice. Eight miRNAs (miR-146a-5p, miR-30c-5p, miR-22-5p, miR-323-3p, miR-361-5p, miR-375-3p, miR-720 and let-7c-5p) were upregulated in the TLR4^−/−^ infected mice. There is clear evidence that the TLR2- and TLR4-mediates the early inflammatory response [88] or pro-inflammatory cytokine production [89,90] and induced inflammation in PD development [22]. Notably, miR-146a-5p upregulated uniquely among the three different infection groups. miR-146a-5p was reported as an inflammation-induced miRNA in PD [44–46].

Several studies reported that miRNAs can intervene in the initiation and modulate the expression of TLRs and multiple components of TLR-signaling pathways such as signaling proteins, transcription factors, and cytokines [91,92]. Activated TLRs induce several TLR-responsive upregulated and downregulated miRNAs [92]. The upregulated miRNA predicts base pairing with sequence in the 3’ UTR of the responsive protein genes and inhibits essential protein expression for TLR activation and affects TLR signaling [93]. Activated TLR2 and TLR4 receptors increased the expression of miR-155, miR-146, miR-147, and miR-9 [92]. Activated TLR4 alone upregulated miR-21, miR-223, miR-125b, let-7e, miR-27b and downregulated miR-125b, let-7i and miR-98. miR-146 has verified TLR-signaling targets of IRAK-1, IRAK-2, TRAF6, in the TLR signaling pathways [92]. miR-155 has verified TLR-signaling targets of MYD88, TAB2, IKKε, and transcription factor targets of FOXP3, C/EBPβ, cytokine targets of TNF, and regulators of SHIP1 and SOCS1 [92].

Based on the above evidence, we interpret that the DE (upregulated and downregulated) miRNAs in wild-type, TLR2^−/−^, and TLR4^−/−^ mice could be responsive miRNAs for activation of TLR2 or TLR4 or together. miR-146a-5p commonly upregulated in wild-type, TLR2^−/−^ and TLR4^−/−^ mice could be a target for TLR signaling in both TLR2/4 receptors. The upregulated miR-15a-5p could target the TLR pathway through TLR4 receptor activation. The ROC curve analysis revealed miR-146a-5p as the most predictable biomarker in PD in the infected mice. The AUC value for 146a-5p was 0.975 in the wild-type mice and 0.950 and 0.940 in the TLR2^−/−^ and TLR4^−/−^ mice, respectively. This positive correlation confirms miR-146a-5p as a polymicrobial-induced PD-inflammatory biomarker among the wild-type, TLR2^−/−^ and TLR4^−/−^ infected mice. Earlier reports suggest that miR-15a-5p modulates the natural killer and CD8^+^ T-cells activation [94] and a vital regulator in inflammation-induced sepsis [95]. miR-15a-5p was reported as an upregulated miRNA in the gingival crevicular fluid of the PD patients [49] and it was also upregulated in the wild-type (AUC:0.870) and TLR2^−/−^ mice (AUC:0.820) with the correlations in AUC values. The present study confirmed that miR-146a-5p and miR-15a-5p are unique TLR2^−/−^ independent PD markers and recommended molecules of periodontal therapy.

In the differentially expressed study, eight miRNAs (miR-22-5p, miR-323-5p, miR-30c-5p, mmu-let-7c-5p, miR-146a-5p, miR-375-3p, miR-361-5p, miR-720) were upregulated in the mandibles of TLR4^−/−^ infected mice. The DE upregulated miRNAs (miR-30c-5p, miR-375-3p, and miR-720) are associated with PD [44,75]. Several of the upregulated miRNAs were associated with systemic diseases such as acute myocardial infarction (miR-22**-**5p) [54,55], coronary heart disease patients (miR-323-3p) [65], and acute coronary syndrome (miR-361-5p) [66]. The studies from Wang et al. also reported that miR-361-5p is an upregulated miRNA with ROC-AUC:0.891 in carotid artery stenosis patients [96]. ROC analysis in the present study confirmed a positive correlation for miR-361-5p (AUC:0.960), miR-375-3p (AUC:0.950), and miR-miR-720 (AUC:0.980) as the most predictive markers for mice infected with periodontitis. miR-361-5p serves as a biomarker to predict acute coronary syndrome [66]. The AUC values for miR-30c-5p and mmu-let-7c-5p were identical (AUC:0.880), and miR-22**-**5p (AUC:0.800), and miR-323-3p (AUC:0.850) has lowest AUC values. miR-22-5p was downregulated commonly in wild-type, and TLR2^−/−^ mice could be a responsive miRNA on TLR signaling through TLR4 activation. The miRNAs of miR-22-5p, miR-361-5p, miR-375-3p, and miR-720 downregulated in TLR2^−/−^ mice were upregulated in TLR4^−/−^ mice. These miRNAs could be TLR-independent signaling markers/ligands. Let-7c-5p in wild-type (downregulated) and in TLR4^−/−^ (upregulated) mice could be a TLR target marker/ligand-activated through TLR2 receptors.

Using the DIANA-miRPath web tool and KEGG analysis, we found that upregulated and downregulated DE miRNAs altered the gene expression in specified pathways further to inflammation, infection, and PD. The KEGG-predicted functional pathway analysis revealed significant findings among the wild-type, TLR2^−/−^, and TLR4^−/−^ mice. The miRNAs upregulated in wild-type, TLR2^−/−^ and TLR4^−/−^ mice have the target pathways of 49, 51, and 67, respectively. The miRNAs upregulated in wild-type and TLR2^−/−^ mice (or) wild-type and TLR4^−/−^ mice have 43 similar target pathways. The upregulated miRNAs in TLR2^−/−^ and TLR4^−/−^ mice have 37 similar target pathways.

The pathways unique for wild-type and TLR2^−/−^ mice, as well as for wild-type and TLR4^−/−^ comparison study, are as follows. Epstein-Barr virus infection, measles, natural killer cell-mediated cytotoxicity, NF-kappa B signaling pathway, tyrosine metabolism, viral carcinogenesis of the TLR2 mice, and adherens junction, ECM-receptor interaction, gap junction, salivary secretion, vascular smooth muscle contraction, etc. of the 30 pathways in TLR4^−/−^ mice. This miRNA expression kinetics was used to identify target genes that can influence the function of PD-related biological pathways. Using the DIANA-miRPath web tool and KEGG analysis, we found that the DE-miRNAs of upregulated and downregulated altered the gene expression in specified pathways further toward inflammation, infection, and PD. The results demonstrated that polymicrobial infection induced PD in wild-type mice with significant ABR and elevated the expression of miR-146a-5p, miR-132-3p, and miR-15a-5p. Polymicrobial infection-induced PD in TLR2^−/−^ and TLR4^−/−^ mice, but none showed significant ABR compared to their sham infection. miR-146a-5p and miR-15a-5p upregulated in the TLR2^−/−^ mice and miR-146a-5p and the other 7 miRNAs upregulated in TLR4^−/−^ mice. miR-146a-5p and miR-15a-5p could be markers for PD in wild-type and TLR2^−/−^ mice. miR-146a-5p, miR-30c-5p, miR-323-3p, and miR-375-3p upregulated from TLR4^−/−^ mice were also reported in human PD, hence considered markers of PD in TLR4^−/−^ mice.

In conclusion, this study explored the expression levels of selected 15 miRNAs that were significantly altered (upregulated/downregulated) during polymicrobial and monobacterial infections in C57BL6/J wild-type, TLR2^−/−^, and TLR4^−/−^ mice, compared to sham-infected controls using RT-qPCR. The data established miR-146a-5p, miR-15a-5p, miR-30c-5p, miR-361-5p, miR-375-3p, miR-720, and mmu-let-7c-5p possibly function as beneficial biomarkers for the therapeutic target for periodontitis. In addition, these data indicate that the above miRNAs may play a critical role (markers/ligands) in the induction of gingival inflammation by regulating TLR2/4 signaling pathways.

## AUTHOR CONTRIBUTIONS

S.J. performed mouse experiments and analyzed the data. A.Y. and J.O. did the molecular analysis of distal organ dissemination; P.G. performed colony PCR for oral swabs; A.R.L.M., E.C., S.R.N., and T.D., performed the Alveolar bone resorption study; J.W. performed data analysis for RT-PCR. I.B. was responsible for histology interpretations. L.K. was responsible for the conception, experimental design, analysis, initial drafting, editing, supervision, project administration, interpretation, and funding acquisition. L.K. and S.J. participated in the final revision and editing of the manuscript. L.K. and E.K.L.C. are the Principal Investigators of the NIH (NIDCR) study. All authors have read and agreed to the published version of the manuscript.

## FUNDING

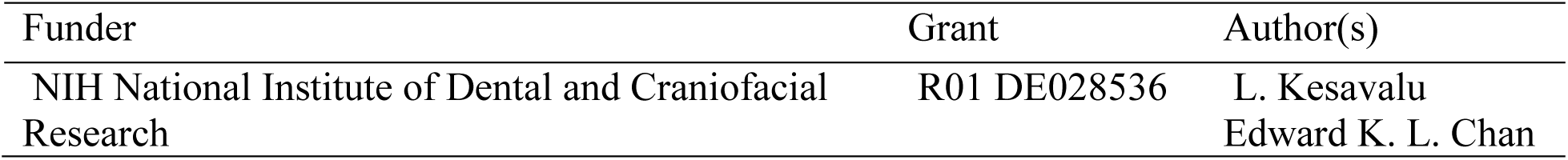

## INSTITUTIONAL REVIEW BOARD STATEMENT

All animal procedures were approved by the University of Florida Institutional Animal Care and Use Committee (IACUC) under protocol number 202200000223.

## INFORMED CONSENT STATEMENT

Not applicable.

